# Targeted ortholog search in unannotated genome assemblies with fDOG-Assembly

**DOI:** 10.1101/2025.09.19.677253

**Authors:** Hannah Muelbaier, Freya Arthen, Vinh Tran, Ina Schaefer, Miklós Bálint, Ingo Ebersberger

**Affiliations:** Applied Bioinformatics Group, Faculty of Biosciences, Goethe University Frankfurt, D-60438 Frankfurt am Main, Germany; LOEWE Centre for Translational Biodiversity Genomics (TBG), D-60325 Frankfurt am Main, Germany; Senckenberg Biodiversity and Climate Research Centre (S-BIK-F), D-60325 Frankfurt am Main, Germany; Institute of Insect Biotechnology, Justus-Liebig University, 35392, Giessen, Germany

## Abstract

Whole genome shotgun sequencing and assembly is routine. However, identifying proteincoding genes in newly assembled genomes remains complex, time-consuming, and labour-intensive. Therefore, most eukaryotic genome assemblies in public databases lack gene annotations reducing their value for evolutionary and functional genomics. Here, we present fDOG-Assembly, a novel tool for targeted, feature architecture-aware ortholog searches directly in unannotated genome assemblies. Benchmarking shows that fDOG-Assembly performs similarly to BUSCO and Compleasm in ortholog identification while offering the advantage of not being restricted to universal single-copy genes. Applied to identify orthologs of 5, 000 human genes in rat and *Nematostella vectensis*, fDOG-Assembly approaches the performance of traditional ortholog search tools that rely on pre-annotated proteomes. Importantly, it can recover orthologs missed by conventional methods because of incomplete gene annotations, helping to fill gaps in phylogenetic profiles. As a case study, we screened 176 soil invertebrate genome assemblies for genes involved in antibacterial compound production. We found that orthologs of β-lactam biosynthesis genes are widespread in springtails, with individual species possessing nearly complete cephamycin biosynthetic gene sets, suggesting they may represent previously unrecognized natural producers of β-lactam antibiotics. Overall, fDOG-Assembly is a powerful resource for orthology-based analyses of the rapidly growing collection of unannotated genome assemblies.

## Introduction

The number and taxonomic diversity of sequenced genomes in public databases is constantly increasing. Continuous advancements in DNA sequencing technologies, combined with near-automated genome assembly workflows further accelerate the rate with which novel assemblies are added (1). This wealth of data can provide unprecedented insights into the evolution and functional diversity of cellular life. However, its value ultimately depends on how comprehensively and accurately the evolutionary relationships between genes encoded in individual species’ genomes and those already represented in databases, particularly those of experimental model organisms, can be determined. The identification of genes whose evolutionary lineages split due to speciation events (orthologs) is commonly used for this purpose (2). Because orthologs minimize the evolutionary time two genes of different species evolve independently, they are also considered the best guess when looking for functionally equivalent proteins across species (3).

A wide range of ortholog search algorithms have been developed, varying in sensitivity, specificity, speed, and scope (4). Comprehensive tools aim to identify orthologs for all genes in a genome. Their computational demands grow rapidly with the number of genes and taxa (5), limiting their scalability. In contrast, targeted tools like fDOG (6), KofamKOALA (7), DeepNOG (8), ODB-mapper (9) and OMAmer (10) extend precomputed ortholog groups using only candidate genes. This facilitates an efficient analysis even of large taxon sets (e.g., (11, 12). Tools like fDOG further assess feature architecture conservation to flag potential annotation errors or functional divergence (13, 14). Crucially, however, all these methods depend on the availability of pre-annotated protein-coding gene sets for the taxa being analyzed. Any gene that was missed during annotation leaves an artificial gap in the presence/absence pattern of orthologs across species resulting in a spurious signal of gene loss. While generating high-quality genome assemblies has become a relatively swift endeavour, their annotation still constitutes a significant bottleneck (1). This is particularly true for eukaryotes with large genomes, where the resulting low gene density paired with long introns and complex gene structures renders an accurate gene annotation a time- and resource-consuming task (15, 16). Accordingly, only 16% (10, 622/64, 534) of the eukaryotic genomes that are currently hosted by the NCBI Genome database, have their genes annotated (NCBI Genome; last accessed 2026/06/10).

Currently, few options are available for a targeted retrieval of orthologs directly from DNA sequence reads or from genome assemblies. Tools like aTRAM (17), HybPiper (18), Kollector (19), Assexon (20) or AliBaSeq (21) have been designed with the intention of compiling large and taxonomically diverse data sets for phylogenomics analyses. These tools use either protein sequences or protein-coding sequences as baits. They then extract matching sequences from contigs that are either assembled on the fly from selected sequence reads or retrieved from existing genome assemblies. Comprehensive annotations of genes or the precise inference of orthology for genes with a history of gene duplications is not the focus of these tools (see (21)). As an alternative, pre-computed orthologous groups (OGs) can be extended with amino acid sequences encoded in transcripts (22–25). The resulting orthology assignments are highly precise. However, only such orthologs are reliably detected across taxa that are consistently highly expressed at the time point of tissue sampling for RNAseq, which can introduce a bias in comparative analyses (e.g., (26)). Only few software solutions exist for identifying orthologs directly in unannotated genome sequences. TOGA (27) utilizes whole genome alignments for projecting gene annotation from a reference to a target genome. The eggNOG-mapper (28) can perform *de novo* gene prediction on raw genomic contigs.

Predicted genes are then assigned to orthologous groups in the eggNOG database when suitable matches are found. BUSCO (23) and Compleasm (29) also annotate protein-coding genes when run in genome mode. Both tools subsequently use profile Hidden Markov Models (pHMMs) and empirically derived bit score thresholds to assign proteins to predefined sets of universal single-copy orthologs.

Tools from this last category can identify orthologs in unannotated genome assemblies, but they come along with several limitations. TOGA, by design, performs exclusively pairwise genome comparisons and relies on whole-genome alignments, making it computationally intensive. In contrast, eggNOG, BUSCO, and Compleasm depend on precomputed, tool-specific collections of orthologous groups and do not support user-defined custom sets. Importantly, none of these tools provide insights into differences in protein feature architecture between reference and target orthologs (13).

Here, we present fDOG-Assembly (fDA), a new software for the targeted identification of orthologs directly from unannotated whole-genome assemblies. fDA integrates orthology inference with an assessment of protein feature architecture similarities between the seed protein and the identified orthologs. We evaluate its performance against established ortholog detection approaches and demonstrate its applicability by screening 176 unannotated genome assemblies of soil-living invertebrates for genes associated with beta-lactam biosynthesis. By enabling ortholog detection without prior gene annotation, fDA expands taxon sampling opportunities and supports consistency-based evolutionary analyses.

## Material and Methods

### Sequence data

For benchmarking the performance of fDOG-Assembly (fDA), we downloaded genome assemblies of 10 metazoan species represented in the QfO reference proteomes (version 2022; Table 1) using the NCBI *genome* package (30). The corresponding annotations of protein-coding genes were retrieved from NCBI or from Ensembl (Table 1). For each assembly, we additionally obtained the cross reference files mapping NCBI to Uniprot ids (gene mapping file) from the QfO reference proteome website ((4) (https://ftp.ebi.ac.uk/pub/databases/reference_proteomes/previous_releases/qfo_release-2022_02_with_updated_UP000000437)). From the same source, we downloaded the reference proteomes of human, mouse and *Drosophila melanogaster.* For each of the three species, we downloaded pairwise orthology assignments against the other 10 metazoan QfO reference proteomes from the QfO benchmark service websites (4) (https://orthology.benchmarkservice.org/proxy/projects/2022/), covering 13 ortholog prediction methods.

**Table 1.** Genome assemblies used in the fDA benchmark.

|  | NCBI<br>Taxonomy<br>ID | Genome Accession | Annotation ID<br>(UniProt) | Assembly<br>source |
| --- | --- | --- | --- | --- |
| *Caenorhabditis elegans* | 6239 | GCA_000002985.3 | UP000001940 | ENA/EMBL |
| *Danio rerio* | 7955 | GCF_000002035.6 | UP000000437 | RefSeq |
| *Drosophila melanogaster* | 7227 | GCA_000001215.4 | UP000000803 | ENA/EMBL |
| *Gallus gallus* | 9031 | GCA_000002315.5 | UP000000539 | Ensembl |
| *Helobdella robusta* | 6412 | GCA_000326865.1 | UP000015101 | Ensembl |
| *Ixodes scapularis* | 6945 | GCA_000208615.1 | UP000001555 | Ensembl |
| *Nematostella vectensis* | 45351 | GCA_000209225.1 | UP000001593 | ENA/EMBL |
| *Rattus norvegicus* | 10116 | GCA_015227675.2 | UP000002494 | Ensembl |
| *Tribolium castaneum* | 7070 | GCA_000002335.3 | UP000007266 | ENA/EMBL |
| *Xenopus tropicalis* | 8364 | GCF_000004195.4 | UP000008143 | RefSeq |

To demonstrate the use of fDA in an applied analysis, we downloaded the genome assemblies of 176 species with a BUSCO completeness over 50% (metazoan_odb10 BUSCO set) from the soil MetaInvert project (31)(NCBI - BioProject ID PRJNA758215).

### Gene sets

We used two sets of orthologous groups for targeted ortholog searches in unannotated genome assemblies. Following the terminology in Ebersberger et al. (2009), we refer to these groups as core orthologs (22). Set 1 comprised the 954 universal single-copy genes of the metazoa_odb10 BUSCO dataset, downloaded from the BUSCO website. For each BUSCO group, we obtained its profile Hidden Markov Model, the block profile, and the majority-rule consensus sequence. The corresponding amino acid sequences were downloaded from the OrthoDB database. To map BUSCO group IDs to UniProt IDs, we used the human representative in each orthologous group, if available, as a query for a *blastp* (v. 2.14.1+) search in the human QfO reference proteome (Table 1). The UniProt ID of the best hit was then mapped to the BUSCO ID if its sequence similarity was at least 90% and the alignment length was at least 70% of the shorter sequence. If the orthologous group did not contain a human sequence, we used the *D. melanogaster* sequence instead. One group lacking both a human and a *D. melanogaster* sequence was ignored. From 954 genes in the *metazoa_odb10* set, we were able to map 917 to a UniProt ID (Supplementary Table S1). For the second set of core orthologs (Set 2), we randomly selected 5000 genes from the human reference proteome.

### Targeted ortholog search

We used BUSCO v5.6.1 and Compleasm v0.2.6 each with default parameter settings. To match the cut-off used by BUSCO, fDA v0.2.0.1 was run with an E-value cut-off of 0.001 for the *blastp* search. Additionally, we invoked the parameter *–gff* to return gene annotations in GFF format. All other parameter settings were kept as default. As the reference species for fDA, we used either human or *D. melanogaster* for Set 1, and human for Set 2.

BUSCO, fDA-tb (tblastn as the search algorithm (32)) and fDA-MP (MiniProt as the search algorithm (33)) were run once with Augustus (34) as the gene predictor and once with MetaEuk (35). fDA-MP was additionally run in fast mode, using directly the MiniProt gene predictions. For all runs invoking Augustus, we adjusted the species-specific model parameters used for the gene prediction to match the respective target species. In the case that no parameter set was provided by Augustus for the target species, we used the one of the closest related species in the Augustus model collection. Ties were broken randomly but we ensured that the same model parameters were used for both fDA and BUSCO. The corresponding mapping file is given as Supplementary Table S2. For the ortholog searches invoking MetaEuk, we used different reference databases for the two sets. For set 1, we used the precomputed reference database provided with *the metazoa_odb10* set in both the BUSCO and fDA runs. For Set 2, we compiled a custom database including all sequences provided by OMA (Release December 2021).

### Core ortholog compilation

For the benchmark set 1, we used the BUSCO groups of metazoa_odb10 as core orthologs for fDA. The helper script fdog.addCoreGroup was used to create the data and formats necessary for fDA from the corresponding multi-fasta files. For the benchmark set 2, we used 5, 000 randomly picked human proteins as seeds and compiled core orthologous groups with fDOG using the species listed in Supplementary Table S3. We set the fDOG parameters *minDist* and *maxDist* to ‘genus’ and ‘kingdom’, respectively. All other values were kept as default. We provide all core orthologs as Supplementary Data S1.

### UniProt ID assignment

The QfO benchmarking framework requires each ortholog to be represented by a UniProt ID. Because fDA, Compleasm, and BUSCO each generate their own gene annotations, we mapped the predicted orthologs to UniProt IDs based on the overlap between their predicted coding exons and the annotated genes in the target species’ genome. Briefly, we extracted the genomic coordinates of the predicted protein-coding exons and compared them with the coordinates of exons in the corresponding genome annotation. We considered a match if two exons reside on the same strand, overlap in their genomic position, and if the cumulative length of all exonic overlaps exceeds 50% of the shorter CDS (Supplementary Fig. S1). If a gene prediction overlapped with more than one reference gene, we selected the gene that maximised the overlap. The reference gene ID in the annotation file was then mapped to UniProt IDs using the ID cross-reference file that is provided together with each of the QfO reference proteomes. The results of the ID cross-referencing are given in Table 2, and the overlap distributions are summarized, for each species, and in the Supplementary Figs. S2-S9.

**Table 2.** Number of inferred orthologs per tested approach.

|  | Gene predictor | # orthologs | #orthologs mapped to UniProt |
| --- | --- | --- | --- |
| fDA-tb<sub>Aug</sub> | Augustus | 9156 | 8273 |
| fDA-MP<sub>Aug</sub> | Augustus | 9457 | 8521 |
| BUSCO | Augustus | 9521 | 8623 |
| fDA-tb<sub>ME</sub> | MetaEuk | 9652 | 8288 |
| fDA-MP<sub>ME</sub> | MetaEuk | 9629 | 8600 |
| BUSCO | MetaEuk | 9815 | 8621 |
| fDA-MP | MiniProt | 9571 | 8684 |
| Compleasm | Miniprot | 9529 | 8743 |

### Running time comparison

Running time measurements were obtained for each benchmark run individually on a compute server equipped with four AMD EPYC 7601 32-core processors. In each run, the 954 genes of Set 1 were searched against 10 genome assemblies. Ten jobs were executed in parallel, with each job allocated 10 CPU cores and 3 GB of RAM per core. For both BUSCO and Compleasm, each job searched for orthologs to all genes in one species. fDA performs the ortholog searches gene by gene. Here, each job searched orthologs for one gene in all species.

### Screening soil-living invertebrates for the presence of the beta-lactam cluster

We screened the unannotated genome assemblies of 176 soil-living invertebrates for genes involved in the biosynthesis of Penicillin G and Cephamycine C (KEGG – map 00311). *Streptomyces clavuligerus* (36) and *Aspergillus nidulans* (37) were selected as reference species for the ortholog search. Seed protein sequences for the two reference species were downloaded from UniProt. Core ortholog groups were computed with fDOG (6) using a species collection containing bacteria and fungi from the RefSeq database release 207 (Supplementary Table S4). Gene names of the seed genes, corresponding reference species, protein IDs used in the reference protein set and UniProt identifiers are given in Supplementary Table S5. The core ortholog groups provided by fDOG served as input for fDOG-Assembly using tblastn and MetaEuk as a gene predictor (fDA-tb_ME_). Phylogenetic profiles were visualised and evaluated with PhyloProfile v.2.0.0 (38).

### Taxonomic assignment

The MetaInvert genome assemblies are likely contaminated with microbial sequences. To distinguish between orthologs that are encoded in the invertebrate genomes, and those that represent microbial contaminations, we performed a taxonomic assignment with DIAMOND v2.1.9 (39). The amino acid sequence of each predicted ortholog was used as query for a DIAMOND search against the NCBI non-redundant database, excluding the species connected to the corresponding assembly. All hits within a bit score margin of 10% of the best hit were considered, and the last common ancestor of the represented species was used as a tentative taxonomic label of the query sequence. For selected genes with inconclusive signals, the entire scaffolds were annotated using MetaEuk with the UniRef90 database as a reference. All identified genes were then taxonomically assigned using the method described above to assess the origin of the orthologs of interest in the context of the full scaffold.

## Algorithm

### Workflow of fDOG-Assembly

fDA performs targeted ortholog detection for a user-defined seed gene in unannotated genome assemblies. The search is guided by a precomputed core ortholog group comprising proteins orthologous to the seed. The workflow includes four main steps: (i) input processing, (ii) identification of candidate genomic regions, (iii) gene prediction within these regions, and (iv) orthology assessment of the predicted proteins (Fig. 1).

**Figure 1.**
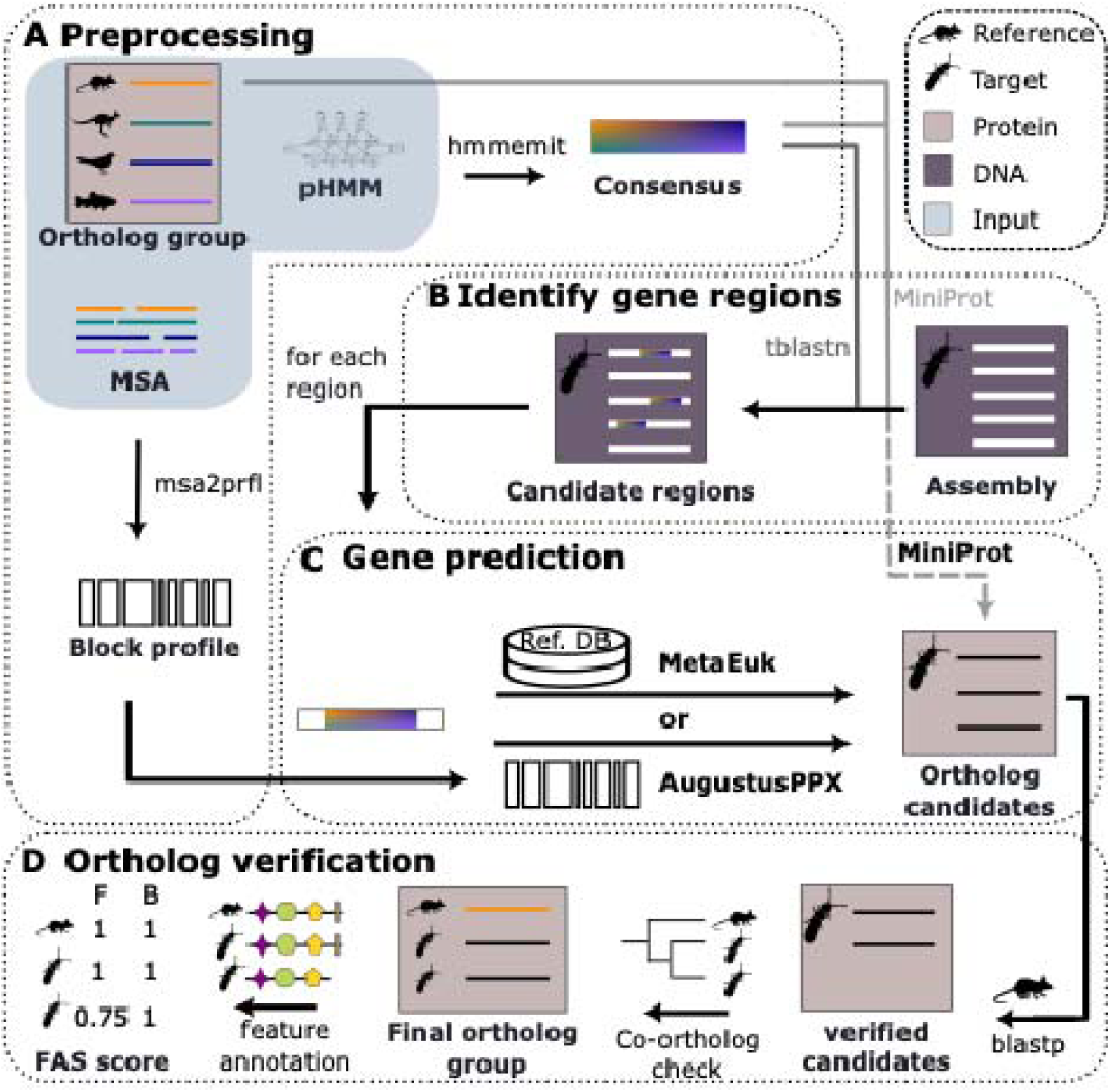
Workflow of fDOG-Assembly. (A) fDA starts its ortholog search from a pre-computed core ortholog group together with a multiple sequence alignment (MSA) of the corresponding amino acid sequences, and a profile Hidden Markov Model (pHMM) trained with the MSA. A consensus sequence is computed from the pHMM, and the MSA is used to produce a block profile. (B) fDA provides two complementary strategies for identifying candidate genomic regions that may contain an ortholog. One strategy uses a tblastn search with the consensus sequence as the query. The alternative strategy employs MiniProt and searches the target genome with both the consensus sequence and all proteins from the core ortholog group. (C) Each candidate region from (B) serves as input for a gene prediction. fDA provides three options for gene prediction, Augustus in combination with the pre-computed block profile, MetaEuk in combination with a reference database or, in the fast mode, MiniProt. (D) To verify the ortholog candidates resulting from (C), the amino acid sequences of the predicted genes are used as queries in a reverse BLASTP search against the protein set of a user-specified reference species. If two or more candidate orthologs are retained, they are accepted as co-orthologs only if their pairwise distance to each other is smaller than their respective distances to the reference protein. Otherwise, the candidate with the smaller distance to the reference is selected. The ortholog search is concluded by assessing pairwise feature architecture similarities between the identified orthologs and the seed gene.

**Figure 2.**
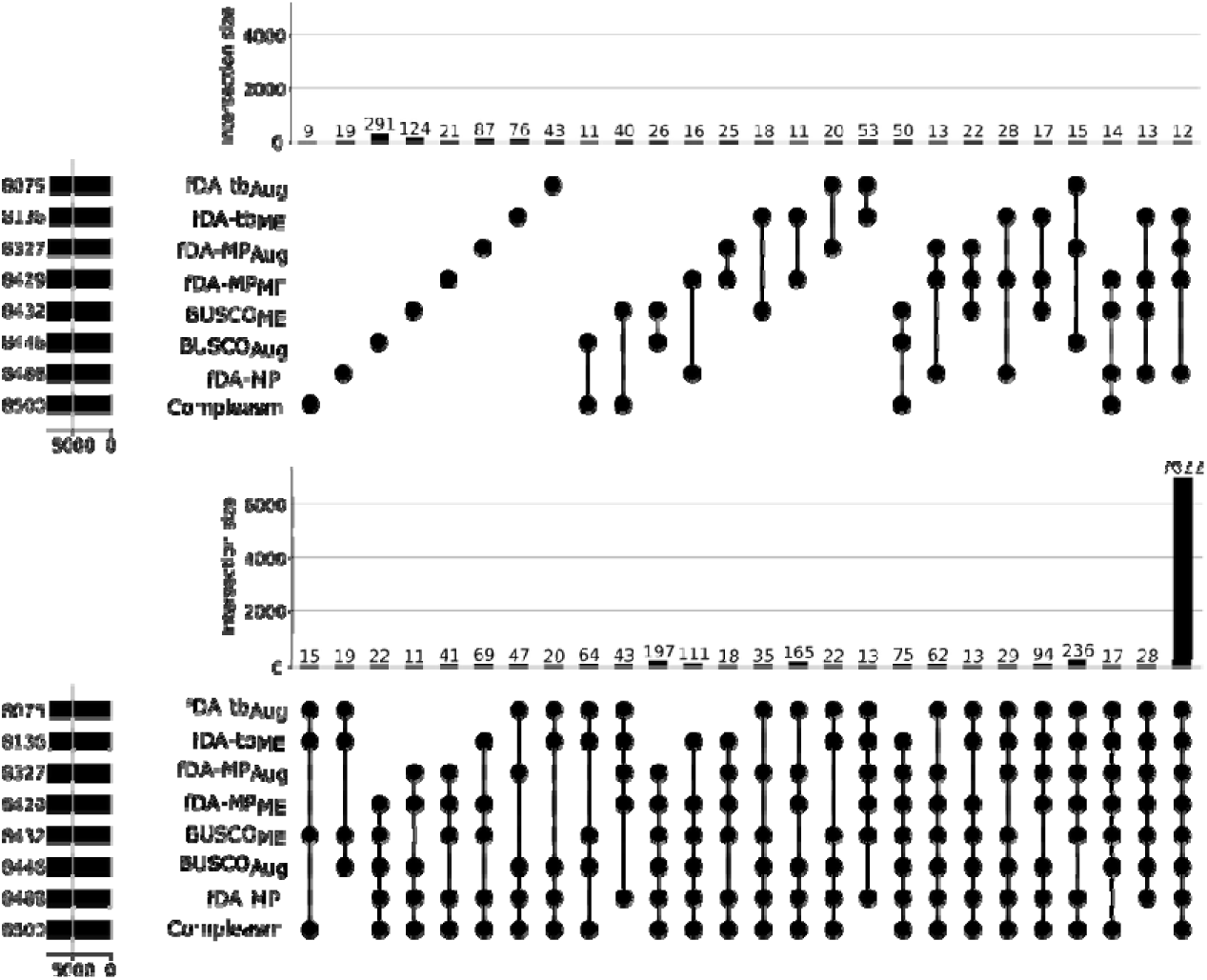
UpSet plot showing the overlap of ortholog assignments made by fDA, Compleasm, and BUSCO. The number of orthologs identified by each individual search is shown in the first column and represented by the horizontal bars. Patterns of stacked filled circles indicate orthologs consistently identified by the corresponding tools, and the total number of orthologs for each pattern is provided above each pattern. Only patterns observed at least 10 times are displayed. The full UpSet plot is provided in Supplementary Fig. S10.

### Input processing

fDA requires a user-provided core ortholog group and the genome sequence of a target species as input (Fig. 1A). Additionally, a reference taxon must be specified that is represented in the core ortholog group. This is optimally the species that is most closely related to the target species, but other choices are possible (see (22)). The sequences in the core ortholog group are aligned with Muscle (40) to form a multiple sequence alignment (MSA), and a profile Hidden Markov model (pHMM) is trained with hmmbuild from the hmmer package (41). Optionally, fDA can use MSAs and pHMMs provided by the user. Next, a majority rule consensus sequence is generated from the pHMM with hmmemit (41). If the user selected Augustus as the gene predictor, a block profile will be computed from the MSA. The purpose of the block profile is to guide the Augustus gene prediction by providing a position-specific frequency matrix that models conserved regions within the MSA. If too many gaps prevent the building of a block profile, gap-rich regions will be pruned from the alignment with *prepareAlign* from the Augustus package, and the block profile building is repeated. If this second attempt fails again, fDA performs the Augustus gene prediction without the block profile.

### Identification of candidate regions

Candidate genomic regions likely to contain orthologs of the seed gene are identified by fDA using either tblastn or MiniProt (Fig. 1B). When using tblastn, the consensus sequence computed in the previous step serves as a query for a search in the target genome. Each hit tags a candidate region for a subsequent gene prediction. Neighbouring regions are recursively joined if the tblastn hits reside on the same DNA strand, and if their distance in the target assembly is below a user-specified maximum (parameter *avIntron*, default value 50, 000bp). By default, up to ten highest scoring regions are then extended on either side (parameter *length extension*, default value 20, 000bp) and the resulting nucleotide sequence is extracted.

When MiniProt is used to identify candidate regions, both the consensus sequence and all sequences from the input core ortholog group are used as queries. If multiple queries generate alignments that overlap the same genomic region, only the alignment with the best-scoring and longest alignment is retained. The ten highest-scoring regions are then selected for ortholog search. Unlike tblastn, MiniProt directly provides a gene prediction as part of the alignment process. In the fast mode of fDA (fDA-MP), this predicted gene model is used directly for ortholog validation. Alternatively, each candidate genomic region can be extended on both sides by a user-defined length (parameter length extension; default: 20, 000 bp), and the corresponding nucleotide sequence is extracted for downstream analysis.

### Gene prediction

For the prediction of genes in the individual candidate regions, fDA offers two options: Augustus or MetaEuk (Fig. 1C). If Augustus was chosen, the user has the option to provide a mapping file assigning each target species a pretrained parameter set. If a block profile is available, it will be used to improve the gene prediction made by the Augustus module *Augustus PPX*. If the choice was MetaEuk, a user-provided database is utilized during gene prediction. If no gene prediction is obtained from Augustus or MetaEuk for a candidate region, fDA uses the MiniProt gene prediction as fallback option provided that MiniProt was used for the initial search in the target genome. Once all genes have been predicted in the candidate regions, the corresponding protein sequences will be propagated to the orthology evaluation step.

### Orthology evaluation

The evaluation of the candidate orthologs resembles the routines in fDOG (6). In brief, each candidate ortholog serves as query for a *blastp* search in the proteome of the reference taxon (Fig. 1D). The candidate ortholog is verified if either of the following conditions are met: (i) the best *blastp* hit is the protein of the reference species that is represented in the core ortholog group, or (ii) the pair-wise Kimura distance between the best *blastp* hit and the reference protein is smaller than or equal to the ortholog candidate and best *blastp* hit. If more than one candidate is verified, we accept them as co-orthologs if their pair-wise distance is smaller than that of either candidate to the reference protein. Otherwise, we select the one with the smaller distance to the reference protein as the ortholog. In the last step of the evaluation, FAS (13) is used to score the similarity of the feature architectures between each of the accepted orthologs and the reference protein. The FAS score ranges from 0, the two proteins share no features, to 1 when their feature architectures are identical. If an ortholog is represented by more than one isoform, as indicated by the corresponding genome annotation file, fDA provides an option to return only the isoform with the highest FAS score to the reference. fDA returns the protein sequences of all identified orthologs, the annotated features and the phylogenetic profile as a tab-delimited file as well as the gene annotation in gff format. The output files can be directly uploaded into PhyloProfile (38) for interactively visualizing and exploring the phylogenetic profiles.

### Helper scripts

fDA provides scripts to automate the preparation of input data as well as the post-process of the output. The scripts *addAssembly* and *addCoreGroup* can be used to set up the directory structure for genome assemblies and precomputed ortholog groups. fDA handles one gene per run. The results of multiple genes can be merged into one phylogenetic profile with the function *mergeOutput*.

### Availability

fDA is available as a part of the fDOG package on GitHub (https://github.com/BIONF/fDOG) under the GPLv3 license, and an accompanying WIKI is available from https://github.com/BIONF/fDOG/wiki/fDOG-Assembly. A set of four Jupyter notebooks provide a step-by-step protocol for performing an fDA analysis with a custom seed sequence, and it is available from https://github.com/BIONF/fDOG/tree/master/examples/fdog_assembly.

## Results

### Benchmark

We benchmarked the performance of fDA using the Quest for Orthologs reference proteomes (release 2022) in combination with the QfO benchmarking service (42). We devised two benchmark scenarios: In scenario 1, we compared the performance of fDA to that of BUSCO when run in ‘genome mode’ and Compleasm. In scenario 2, we searched for orthologs to 5, 000 randomly selected human genes in two species. By their design, BUSCO and Compleasm cannot use an external dataset with multi-copies orthologs. Therefore, we assessed the performance of fDA by comparing its orthology assignments to those made by 13 standard ortholog search tools that utilize proteomes (Supplementary Table S6).

### Scenario 1 - Comparison of fDA against BUSCO and Compleasm

We benchmarked the three tools using 954 conserved genes from the BUSCO metazoa_odb10 dataset. Orthologs were inferred in 10 species selected to represent the diversity of Metazoa (Supplementary Table S2). BUSCO benchmarks were performed using both Augustus and MetaEuk as gene predictors. fDA was evaluated in five search modes that differ in candidate region identification and gene prediction. fDA-tb_ME_ and fDA-tb_Aug_ use tblastn to identify candidate regions, followed by gene prediction with MetaEuk and Augustus, respectively. fDA-MP_ME_ and fDA-MP_Aug_ use MiniProt for candidate region identification and MetaEuk or Augustus for gene prediction. The fifth mode, fDA-MP, is the fastest configuration. It relies exclusively on MiniProt for both candidate region identification and gene prediction. The total number of inferred orthologs across all species is shown in Table 2, and the number per species is given in Supplementary Table S7. Comparing only runs using the same gene predictor, BUSCO identified slightly more orthologs compared to fDA. Both tools identified consistently more orthologs when using MetaEuk instead of Augustus. fDA-MP and Compleasm, which both rely only on MiniProt, perform comparably.

### Performance of fDA, BUSCO and Compleasm compared to proteome-based tools

We used the QfO benchmarking service to compare fDA, BUSCO, and Compleasm among each other and with 13 proteome-based ortholog prediction tools (Figure 3). Differences among the assembly-based methods were modest, but BUSCO_Aug_ performed slightly worse than the other assembly-based approaches across the three benchmark challenges. Overall, assembly-based methods showed similar precision compared to the proteome-based tools but had lower sensitivities. To better characterize this difference, we focused on a reference set of 5, 669 ortholog pairs consistently identified by all 13 proteome-based tools (Table 3). Within this reference set, Compleasm and fDA-MP missed the fewest orthologs, whereas fDA-tb_ME_ missed the most. Lastly, we assessed the consistency between orthology assignments made by genome-based tools and those made by the proteome-based tools (Fig. 3B). BUSCO_Aug_ and fDA-MP_Aug_ resulted in the highest number of assignments that were not reproduced by any of the proteome-based tools.

**Figure 3.**
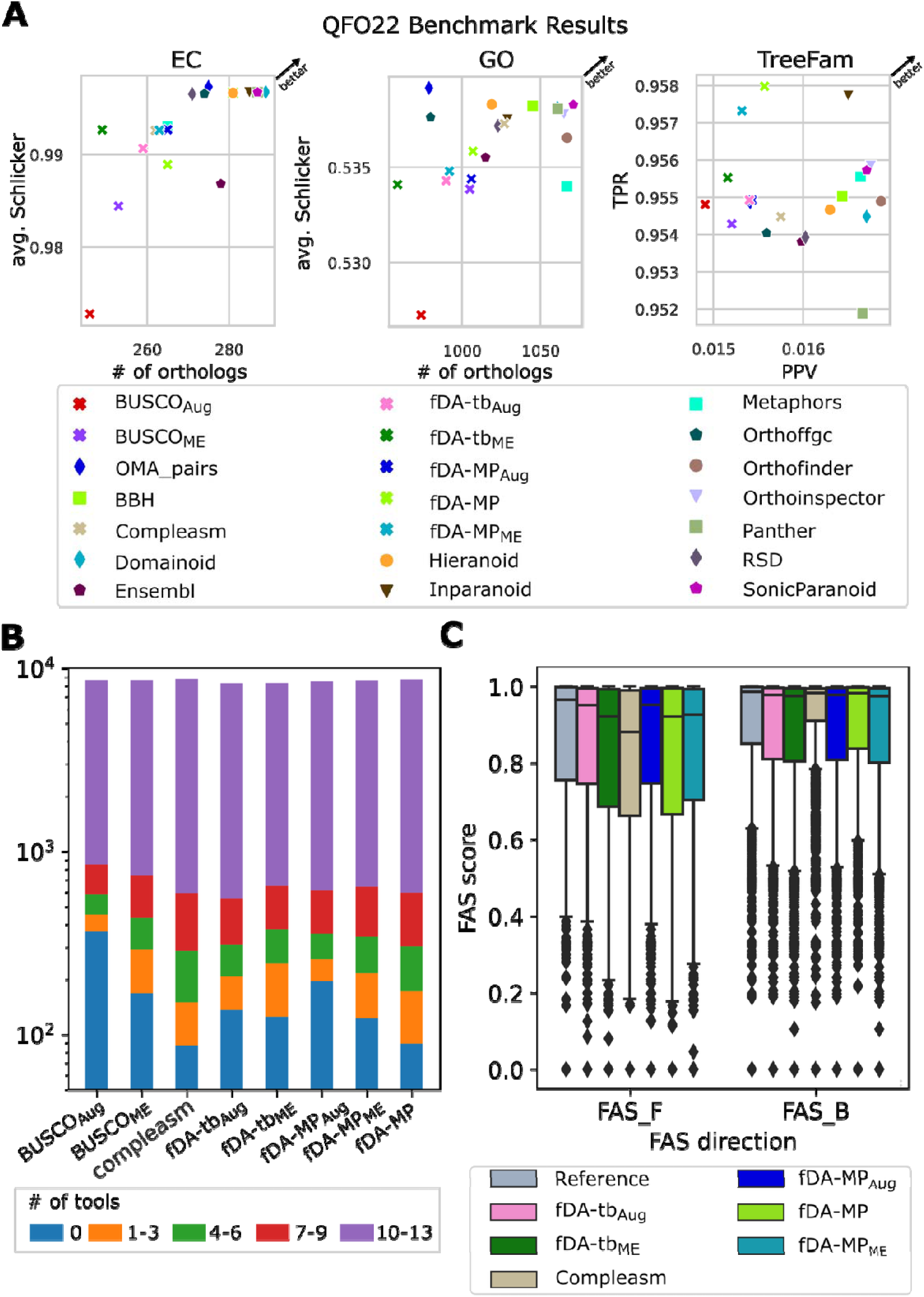
Benchmark results Scenario 1. A) Performance of fDA, BUSCO, and Compleasm in identifying orthologs to the metazoa_odb10 set in the genome assemblies of 10 metazoan species. Performance metrics of the proteome-based ortholog search tools are given for comparison. EC – average semantic similarity (Schlicker score) of the EC annotation between ortholog pairs; GO – average semantic similarity (Schlicker score) of the GO annotation between ortholog pairs; TreeFam – Agreement with curated reference gene phylogenies provided by TreeFam-A. PPV – positive predictive value; TPR – True positive rate. For details see (43). B) Consistency of orthology assignments from assembly-based tools with those from proteome-based tools. The color code indicates how many proteome-based tools support each assignment made by the respective assembly-based tool. Bar heights specify the number of ortholog assignments within each category (log scale). The plot using linear scaling is provided as Supplementary Fig. S11 C) Distributions of pair-wise feature architecture similarity (FAS) scores for orthology assignments using pre-annotated proteomes (Reference) compared to fDA and Compleasm. FAS_F – Feature architecture similarity score using the seed protein as reference; FAS_B – FAS score using the ortholog as reference (see (13)). tb – tblastn; MP – MiniProt; Aug – Augustus; ME – Metaeuk.

**Table 3.** Missed consensus ortholog pairs across assembly-based orthology inference methods (reference set: n = 5, 669) Subsequently, we mapped orthologs detected in the genome assemblies of the ten target species to UniProt IDs (Table 2, Supplementary Fig. S1, and Supplementary Figs. S2-S9). For approximately 10% of orthologs, no UniProt ID could be assigned. These were excluded from downstream analyses. With UniProt IDs available, we directly compared the results of fDA, BUSCO, and Compleasm (Fig. 2 and Supplementary Fig. S10). Most orthologs (7, 022) were identified by all tools, independent of the search strategy. An additional 1, 170 orthologs were detected by at least six of the eight approaches. Fifty ortholog pairs were recovered by Compleasm and both BUSCO runs, but not by fDA. The number of orthologs inferred only by one search strategy is highest for BUSCO_Aug_ (291) and lowest for Compleasm (9). Thus, the orthology assignments are highly consistent across tools, although each method differs in the assignment of a small subset of orthologs.

|  | Missed | Fraction |
| --- | --- | --- |
| **fDA-tb<sub>Aug</sub>** | 311 | 5.49% |
| **fDA-tb<sub>ME</sub>** | 380 | 6.7% |
| **fDA-MP<sub>Aug</sub>** | 199 | 3.51% |
| **fDA-MP<sub>ME</sub>** | 196 | 3.46% |
| **fDA-MP** | 128 | 2.26% |
| **BUSCO<sub>Aug</sub>** | 281 | 4.96% |
| **BUSCO<sub>ME</sub>** | 267 | 4.71% |
| **Compleasm** | 91 | 1.61% |

### Genes exclusively detected by the assembly-based tools close gaps in phylogenetic profiles

So far, the benchmark scenario 1 assigns fDA, but also BUSCO and Compleasm, a very good specificity and only a slightly lower sensitivity compared to the proteome-based tools. We next asked whether the assembly-based tools can identify orthologs that have been missed during genome annotation. To investigate this, we examined all orthology assignments to BUSCO genes that were not confirmed by any proteome-based tool. This builds on the assumption that genes in BUSCO sets are rarely lost during evolution. Thus, gaps in the phylogenetic profiles of genes in the metazoa_odb10 dataset either indicate rare gene loss or annotation errors.

In metazoa_odb10, we found 214 cases in which protein-based tools identified orthologs in 9 of the 10 target species (Supplementary Table S8). We then assessed whether orthologs uniquely detected by the assembly-based methods in the tenth species could fill these gaps. (Table 4). To distinguish between likely spurious and plausible orthologs, we applied the following logic: if a UniProt ID could be assigned to the detected ortholog, it probably reflects a false positive, since the same ortholog should have been found by proteome-based tools. If no UniProt ID could be assigned, it may indicate a true ortholog missed during annotation.

**Table 4.**
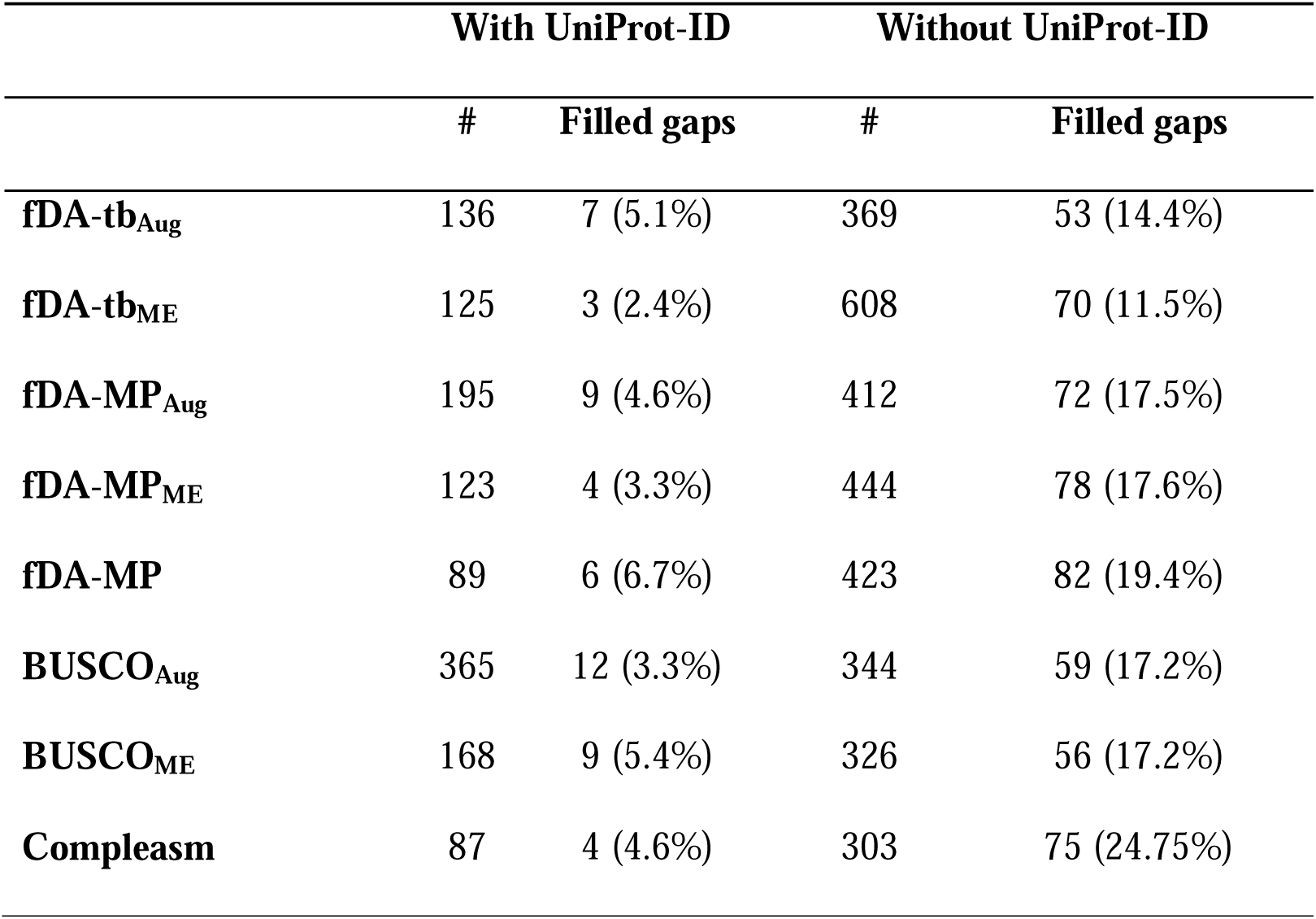
Orthology relationships identified by assembly-based approaches that are not supported by protein-based methods. Total number of orthology assignments is given in Table 2.

Among orthologs with an assigned UniProt ID, only a few filled phylogenetic gaps (2.4– 6.7%, depending on the method). In contrast, orthologs without a UniProt ID filled gaps more frequently (11.5–24.75%). This supports the view that targeted searches in unannotated genomes can indeed recover genes overlooked in standard annotations (see Supplementary Figure S12 for an example).

### Feature architecture comparisons between seed proteins and inferred orthologs

Thus far, we have focused on the performance of fDA in identifying orthologs in unannotated genomes. We next examined how the different fDA search modes affect protein feature architecture similarities between detected orthologs and reference proteins. As a best-case benchmark, we used the distribution of FAS scores derived from ortholog pairs based on the pre-annotated proteins in the QfO reference proteomes. All other results were compared against this benchmark. We included Compleasm in this comparison because it relies solely on MiniProt and does not perform additional gene prediction. BUSCO was not included, as its combination of search strategy and gene prediction is covered by fDA. For this analysis, we used a set of 5, 075 proteins for which all 13 proteome-based tools, all fDA search modes, and Compleasm consistently identified an ortholog.

For each ortholog pair, we computed pairwise FAS scores and plotted the resulting distributions by method (Fig. 3C). fDA with Augustus as gene predictor produced FAS_F score distributions that closely matched the best-case benchmark. The use of MetaEuk reduced the median FAS_F score. The lowest values were observed for MiniProt-only approaches (fDA-MP and Compleasm). A decrease in FAS_F indicates loss of protein features in the predicted ortholog. This suggests that Augustus generates gene structures most consistent with the QfO reference annotations. MetaEuk, and more strongly MiniProt, tend to miss exons during gene prediction. This leads to truncated feature architectures in the predicted proteins.

The patterns differ for FAS_B scores, which decrease when additional features are present in the ortholog that are absent in the reference protein. Here, differences between methods are minor, and median FAS_B scores are largely similar. This suggests no strong tendency for any gene predictor to introduce spurious exons or artificially extend feature architectures.

However, Compleasm shows a narrower FAS_B distribution than the best-case benchmark. This is consistent with the FAS_F results and indicates a tendency toward truncated feature architectures.

### Running time

For benchmarking the runtime of fDA, BUSCO, and Compleasm, we used the same computational resources to search for orthologs to 954 genes (Set 1) in 10 metazoan species. Compleasm was the fastest method with 1h 21 min followed by fDA-MP (2h 28 min). Adding the Augustus gene prediction doubled the runtime (fDA-MP_Aug_: 4h 59) and replacing MiniProt by tblastn prolonged the runtime by almost an hour (fDA-tb_Aug_: 5h 51 min).

BUSCO_ME_ needed 6h 3 min and 45 seconds, fDA-MP_ME_ 6h 20 min and 35 sec, and BUSCO_Aug_ 6h 32 min. fDA-tb_ME_ was with 11h 27 min the slowest search mode. The running time differences between BUSCO_ME_ and fDA-tb_ME_ is explained by the fact that fDA invokes MetaEuk separately for each gene, whereas BUSCO performs a single MetaEuk analysis for the entire genome assembly. The mean running time per gene across 10 species was 1 min 29 seconds for fDA-MP, 2 min 37 seconds for fDA-MP_Aug_, 3 min 23 seconds for fDA-MP_ME_, 3 min 34 seconds for fDA-tb_Aug_ and 7 minutes and 9 seconds using fDA-tb_ME_.

### Benchmark Scenario 2

Thus far, the benchmark was limited to universal single-copy genes, as they are a prerequisite for running BUSCO and Compleasm. Because fDA does not have this requirement, we next extended the benchmark to identify orthologs for 5, 000 randomly selected human proteins in two target species. We chose rat (*Rattus norvegicus*) as a representative of species that is more closely related to humans (median time to most recent common ancestor: 87 MY (44)). The cnidarian *Nematostella vectensis* served as an example for an evolutionarily more distantly related species (median time to most recent common ancestor: 715 MY (44)) where the higher sequence divergence of protein coding sequences is likely to render the identification of orthologs in unannotated genomes harder (45). In brief, we computed core orthologous groups with fDOG (Supplementary Data S1) using a subset of the QfO22 reference species excluding both rat and *N. vectensis* (Supplementary Table S3) and the human protein as the seed. Subsequently, we extended these core orthologous groups with further orthologs in the genome sequences of the two target species. The results were processed as described for Scenario 1, and the resulting ortholog assignments with an UniProt-ID are summarized in Supplementary Table S9.

Fig. 4 provides a comparison between these results and those obtained with conventional ortholog search tools using the pre-annotated protein sets for the two target species. Similar to the Benchmark Scenario 1, fDA performs comparable to the proteome-based in both benchmark tests (EC and GO), but it has a slightly lower sensitivity. From a set of consensus orthologs that were identified by all proteome-based tools (3, 550 orthologs), fDA-tb_Aug_ identified 87, 2% and fDA-tb_ME_ identified 90.6%. fDA-MP_Aug_ found 89.2%, fDA-MP_ME_ 92.9% and fDA-MP found with 94.6% the highest fraction of the consensus pairs.

**Figure 4.**
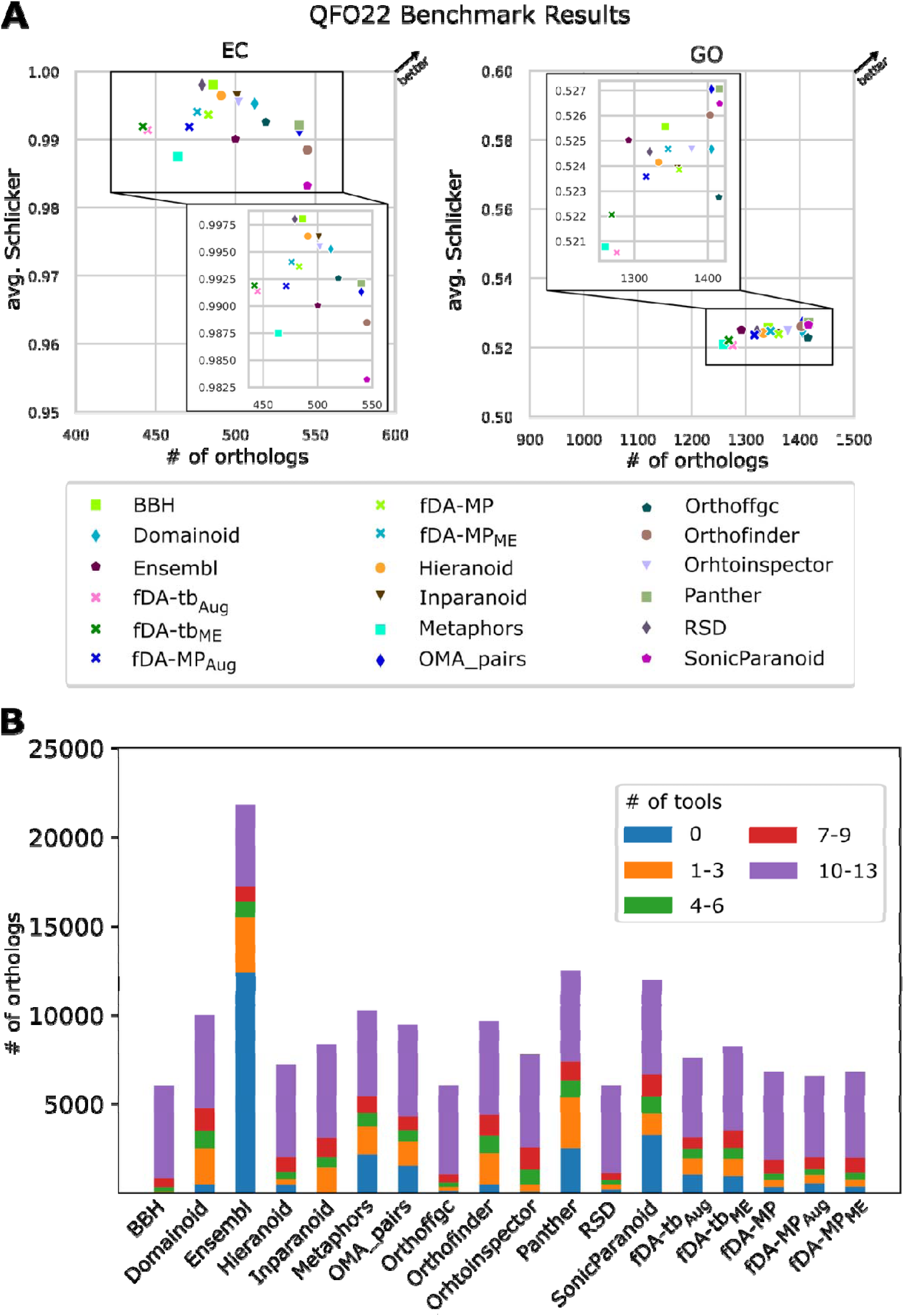
Benchmark results Scenario 2. A) Performance of fDOG-Assembly in identifying orthologs to 5, 000 human proteins in *Rattus norvegicus* and *Nematostella vectensis*.

Compared to benchmark scenario 1, the fraction of missed consensus orthologs is higher (see table 3), which could be due to the added complexity introduced by the presence of co-orthologs. We therefore assessed the performance of fDA in identifying orthologs to genes with a gene duplication history. We classified the orthologous groups generated by InParanoid, a high-performing pairwise orthology inference tool, into 1:1, 1:n, n:1, and n:n categories, where the numbers indicate the number of co-orthologs present in each of the two species being compared. We then assessed the proportion of ortholog pairs induced from these groups that were recovered by fDA with the various search modes (Table 5 (rat) and Supplementary Table S10 (*N. vectensis*). As expected, fDA performs best on 1:1 groups, where it recovers up to 97% of the groups (3873/4010). For n:1 groups, it still recovers 88% of the groups (24 of 27) and 85% of the co-orthologous relationships (54 of 63). n:n groups provide the biggest challenge. fDA recovers up to 19 of the 22 groups with up to 45% of the co-orthologs (156 of 348). Lastly, we determined, for each tool, how many of its orthology assignments were consistent with those of other tools (Fig. 4B). This revealed that most orthologs that were identified with fDA were also found with other tools.

**Table 5.** Performance of fDA in recovering human – rat co-orthologous relationships.

| **Group layout<sup>s</sup>** | **1:1** | **1:n** | **n:1** | **n:n** |
| --- | --- | --- | --- | --- |
| **Orthology assignments: #groups (#pairs)** |  |  |  |  |
| **InParanoid** | 4010 (4010) | 114 (379) | 27 (63) | 22 (348) |
| **fDA-tb<sub>Aug</sub>** | 3661 (3661) | 96 (205) | 21 (45) | 19 (156) |
| **fDA-tb<sub>ME</sub>** | 3673 (3673) | 97 (207) | 19 (38) | 17 (89) |
| **fDA-MP<sub>Aug</sub>** | 3743 (3743) | 94 (169) | 24 (50) | 19 (87) |
| **fDA-MP<sub>ME</sub>** | 3829 (3829) | 97 (167) | 24 (54) | 19 (80) |
| **fDA-MP** | 3873 (3873) | 99 (177) | 23 (50) | 19 (88) |

<sup>s</sup> number of human co-orthologs : number of rat co-orthologs

In summary, our benchmarks reveal that fDOG-Assembly is a powerful tool to identify orthologs in unannotated eukaryotic genome sequences. Contrary to BUSCO and Compleasm, it does not require genes to be single copy. This allows to use any gene as input for a fDOG-Assembly search, which substantially extends the application spectrum compared to BUSCO and Compleasm. Moreover, it also compares the feature architectures of the identified ortholog, tracing both incompletely identified genes and evolutionary changes of the feature architecture.

In the following, we demonstrate the application of fDA by tracing a biosynthetic pathway for β-lactam biosynthesis in unannotated genomes of soil invertebrates.

Performance metrics of the proteome-based ortholog search tools on the same challenge are given for comparison. EC – average semantic similarity of the EC annotation between ortholog pairs; GO – average semantic similarity of the GO annotation between ortholog pairs. For details see (43). The inlays provide a zoomed view on the individual results. B) Consistency of assignments across ortholog search tools. The color code indicates the number of tools that support an orthology assignment by the respective method. Bar heights specify the number of ortholog assignments within each category.

### Beta-lactam gene cluster in soil-living invertebrates

The β-lactam biosynthetic pathway comprises a series of enzymatic reactions that lead to the production of β-lactam compounds, many of which exhibit antimicrobial activity (46). Initial evidence exists that some springtails (Collembola; Hexapoda) can biosynthesize beta-lactams (47). However, Suring and colleagues investigated only 22 springtails alongside with 16 other apterygote species related to springtails (Protura and Diplura). Of these species, 36 were only represented by transcriptomes, which results in the missing of any gene that is not sufficiently highly expressed under the sampled conditions. Therefore, it is still largely unclear how widespread the presence of beta-lactams is within springtails, and whether other soil living invertebrates might also be able to synthesize them. To shed further light on this issue, we used fDA to trace orthologs to the enzymes comprising the biosynthesis pathways for Penicillin G of *Aspergillus nidulans* and Cephamycin C of *Streptomyces clavuligerus* (see Fig. 5A) in the genome assemblies of 176 soil-living invertebrates (31). Species names and corresponding BUSCO values are given in Supplementary Table S11. The resulting phylogenetic profiles are given as Supplementary Data S2. Fig. 5B shows a concise 3D UMAP representation of the data (see (6)), and the full taxon-gene matrix is given as Supplementary Fig. S13. We detected orthologs of at least one enzyme in the genome assemblies of 152 invertebrates. The UMAP reveals that the taxa can be categorised into six groups, each characterized by a specific enzyme signature (Fig. 5C). Notably, three clusters (C3, C4, and C6) comprise taxa in which fDA detected orthologs of both pcbAB and pcbC, the two enzymes synthesizing the β-lactam ring. Taxa in C5 harbour orthologs of pcbAB and cefD, but pcbC seems absent. Interestingly, C4 and C6 are taxonomically homogeneous and comprise mainly springtails, except for two mites in C4 and a tardigrade in C6 (*Isohypsibius dastychi*). Clusters 3 and 5 are a mix of both springtails and mites (Arachnida), where Cluster 3 additionally comprises a single nematode, *Acrobeloides thronei (*see Supplementary Table S12 for the precise cluster compositions).

**Figure 5.**
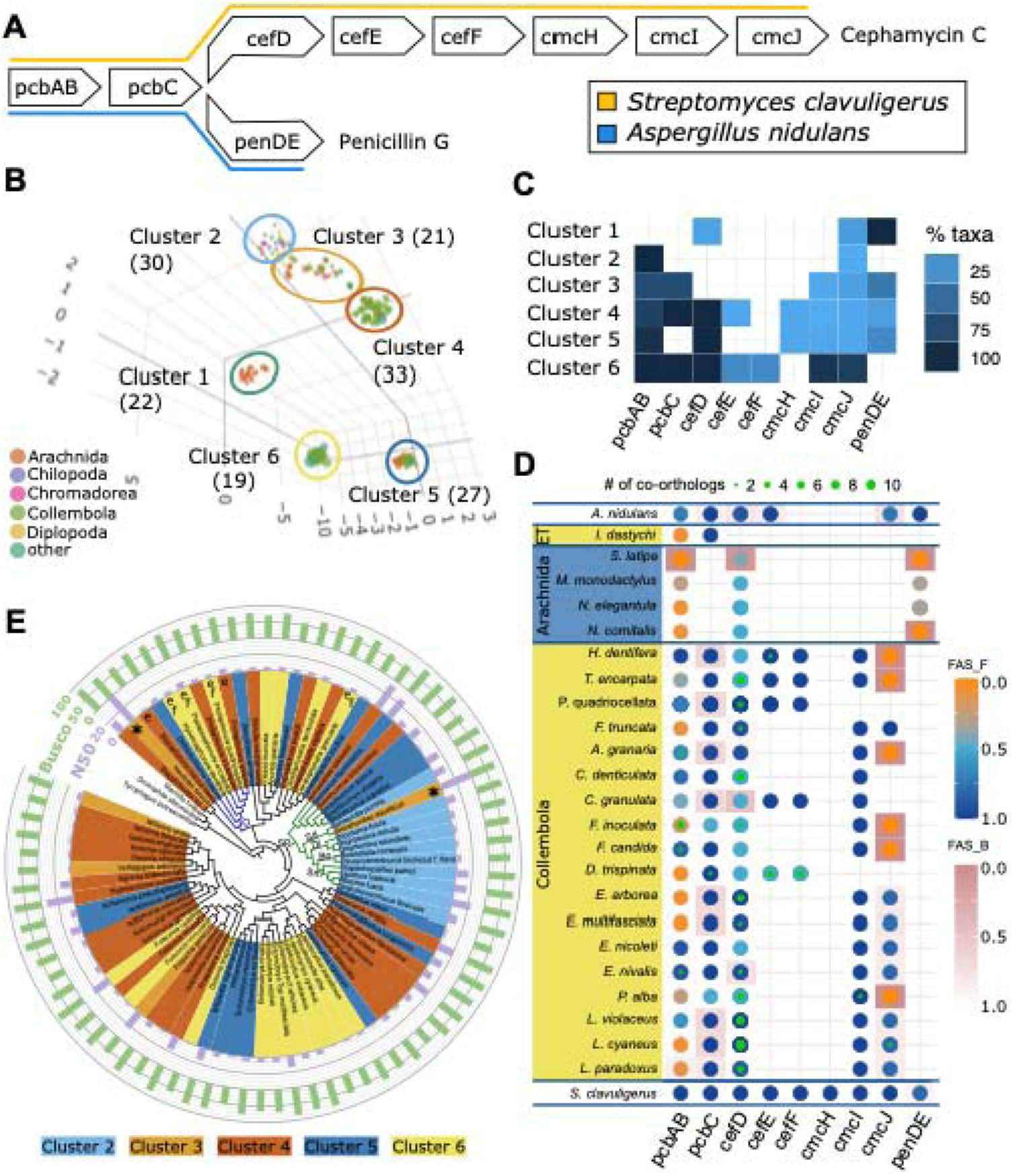
Abundance of orthologs to β-lactam synthesizing genes in the genome assemblies of soil invertebrates. A) Biosynthetic pathways to produce Cephamycin N (*Streptomyces clavuligerus;* bacteria) and Penicillin G (*Aspergillus nidulans;* fungi). B) UMAP ordination representing for each investigated soil invertebrate the profile of orthologs to β-lactam synthesizing enzymes. The corresponding taxon – gene matrix is shown in Supplementary Fig. S13). Dots represent species, and the dot colour informs about the corresponding taxonomic class. Cluster assignment was done by eye. C) The heat map shows the relative abundance of taxa with an ortholog of the indicated gene in its genome assembly for each cluster. D) Taxon gene matrix for selected members of clusters 5 (blue) and 6 (yellow). *A. nidulans* and *S. clavuligerus* are the seed taxa used in the ortholog search. A dot indicates that an ortholog to the corresponding seed gene could be identified in an invertebrate genome assembly. A green inlay indicates that two or more co-orthologs were detected, and the diameter of the inlay informs about the number of co-orthologs. Dot colour represents the protein feature architecture similarity (FAS) score between the seed and the ortholog, using the seed protein as reference. A FAS score of 1 (blue) indicates that the two proteins share the same feature architecture, whereas a score of 0 (orange) indicates no shared features. The cell colour informs about the FAS score using the ortholog as a reference. If more than one ortholog was detected, the one with the highest FAS score was selected for display. ET – Eutardigrada. E) Phylogenetic distribution of β-lactam signatures in springtails. Species in the phylogeny are coloured according to their cluster assignment (see B). The inner ring depicts the N50 sizes of the corresponding genome assemblies in kilo bases (kb). Assemblies with an N50 above 40 kb are indicated with an ‘*’. *Neelides folsomi* – N50: 149, 788bp; *Sminthurides aquaticus* – N50: 60, 729bp. The outer ring gives the BUSCO completeness score (Busco set *metazoa_odb10*). ‘e’ and ‘f’ indicate the presence of orthologs to cefE and cefF, respectively. Phylogenetic tree topology is taken from (12). Blue clade: Onychiurinae; Green clade: Symphypleona. Taxon abbreviations: O – Onychiurinae; SP – Symphypleona; Sd - Sminthurididae; Di – Dicyrtomidae; Bo – Bourletiellidae; Sm – Sminthuridae.

### β-lactam biosynthetic enzymes - intrinsic genes or microbial contaminations

The presence of β-lactam synthesizing enzymes in springtails has been proposed before (47). Mites, tardigrades and nematodes have, to our knowledge, not been reported to produce β-lactams, thus far. However, many genome assemblies, especially of taxa where the small body size requires the entire organism for DNA extraction, are ridden with microbial contaminations. Because ortholog search tools cannot differentiate between intrinsic genes encoded in the invertebrate genomes and those that stem from contaminating sequences, we performed a DIAMOND taxonomic assignment for all orthologous sequences detected in the species highlighted in Fig 5D. We focused on clusters 5 and 6 because the enzyme composition differs from what was described earlier for springtails (see (47)). Genes were flagged as native to the respective target organism if they were assigned to an invertebrate.

They were identified as contaminations if their taxonomic assignment was within bacteria or fungi (Supplementary Table S13). We removed these from the phylogenetic profiles and did not consider them further in the analysis. The origin of genes was flagged as inconclusive if the taxon set in the DIAMOND hit list predominantly consisted of bacterial or fungal taxa and only a single hit to an invertebrate was listed. Due to the isolated identification of orthologs of cefE and cefF and an inconclusive evaluation of their origin, the full scaffolds harboring these genes were evaluated. In total, 14 of all 158 reviewed genes appear as microbial contaminations and for further 20 genes the signal remained inconclusive. In the subsequent analyses, we did not consider these genes, and they are not represented in Figs. 5D and 5E. For cefE and cefF, strong evidence supporting a native origin was obtained from analyses of neighboring genes in four of the five species examined (see Supplementary Table S14). Thus, although microbial contamination is present in the genome assemblies of soil invertebrates, it is unlikely to account for the detected β-lactam biosynthesis genes. Instead, these genes appear to be widespread in springtails and occur sporadically in mites and other soil-dwelling invertebrates.

While orthologs of genes involved in β-lactam biosynthesis are present in many genomes, their functional relevance remains uncertain. For example, both springtails and mites in cluster 5 encode pcbAB and cefD, but lack pcbC, which acts upstream of cefD in the β-lactam biosynthesis pathway (see Fig. 5A). This incomplete gene set raises the question of whether in these cases pcbAB and cefD together participate in a functional biosynthetic pathway.

Taxa in cluster 6 contain the richest enzyme sets observed in springtails thus far. Individual species comprise orthologs to seven of the nine investigated beta-lactam synthesizing enzymes (see Fig. 5D). Except for cmcH, they seem to feature the full biosynthetic pathway for Cephamycin C. Note that cefE of two species (*T. encarpata* and *C. granulata*) could be taxonomically assigned only to the Opisthokonta (see Supplementary Table S13), but the respective genes are embedded in genomic contexts that can be unambiguously assigned to the springtails (Supplementary Table S14). This provides strong evidence that the enzymes are indeed encoded by intrinsic genes making it likely that these taxa can produce a substance very similar to Cephalomycin C.

### Phylogenetic distribution of beta-lactam biosynthesis

The analysis, thus far, has revealed that beta-lactam synthesizing enzymes are prevalent in springtails. Individual species seem to possess almost the entire biosynthetic pathway necessary for producing Cephalomycin C, or at least a similar substance. To put the analysis into an evolutionary context, we projected the cluster assignments of the individual species (Fig. 5B) to the springtail phylogeny (Fig. 5E). We complemented this data with information about genome assembly completeness and assembly contiguity. Taxa in cluster 4, which is characterized by the presence of orthologs to pcbAB, pcbC and cefD, span the full diversity of the springtails. This suggests that the corresponding genes had been acquired already by the last common ancestor of springtails. Members of cluster 6 are spread across the tree yet with a tendency of accumulating in individual clades. This is consistent with a multiple and independent introduction of cmcI and cmcJ during springtail diversification. However, an early acquisition followed by multiple losses cannot be ruled out. Notably, three of the four species with the largest genetic repertoire encoding β-lactam synthesizing enzymes (cluster 6) are closely related and form a clade within the Onychiurinae (blue clade in Fig. 5E). The earliest diverging species in this clade, *Mesaphorura macrochaeta*, also harbours multiple co-orthologs to cefE. Thus, cefE and cefF likely have been acquired already by the last common ancestor shared by all members of this clade. Unfortunately, the quality of the available assemblies is low overall, with N50 values often in the range of 20 kb or below, and BUSCO completeness scores rarely above 75% (See Fig. 5E and Supplementary Table S11). In cases where an ortholog was not detected in an individual species, it is virtually impossible to differentiate whether a gene is truly absent, or its lack is a technical artefact. Yet, the absence of orthologs to β-lactam-synthesizing enzymes in the species subsumed in cluster 2 is a remarkable exception. Within the Symphypleona (green clade in Fig. 5E), there exists a monophyletic clade of four families (*Sminthurididae, Dicyrtomidae, Bourletiellidae, and Sminthuridae*) that consistently lack, with very few exceptions, all β-lactam synthesizing enzymes except for pcbAB. This suggests that pcbC and cefD were lost during the diversification of *Symphypleona* and after the families diverged from their last common ancestor shared with earlier branching members within the Symphypleona that belong to clusters 4 or 5.

## Discussion

Orthologs are essential for tracing the evolutionary trajectory of genes and their functions across species. The resolution of ortholog-based analyses improves with the number of taxa included in the search and with the integration of accessory data that aids in assessing functional diversification between orthologs (13). Large-scale biodiversity genomics initiatives are increasingly fueling such analyses (31, 48–50). However, most publicly available genome assemblies are either unannotated or are associated with only rudimentary gene predictions. This excludes many genomes from orthology analyses reducing the taxonomic resolution of individual studies.

fDA was developed to address this challenge by extending the currently limited toolkit for ortholog identification in unannotated genomes. Unlike existing approaches, fDA integrates targeted gene annotation, robust orthology inference, and comparative analysis of protein feature architectures into a single workflow. This combination not only facilitates ortholog detection but also provides an additional layer of quality assessment by comparing the feature architectures of predicted orthologs to those of the seed protein.

fDA offers five search modes that balance runtime, sensitivity, and gene model accuracy. The MiniProt-based fast mode achieves the shortest runtimes while maintaining high sensitivity. However, this speed advantage comes at the cost of less complete gene models, as reflected by lower feature architecture similarity (FAS) scores between seed proteins and predicted orthologs (see Fig. 3). In contrast, when Augustus is used for gene prediction, the sensitivity slightly decreases, but FAS score distributions closely match those obtained from pre-annotated reference proteomes, indicating improved gene model quality. Overall, we find that the choice of gene predictor has a stronger influence on the performance of fDA than the method used to identify candidate genomic regions. Nevertheless, tblastn seems to slightly perform better than MiniProt in recovering co-orthologs from more distantly related species (see Supplementary Table S10).

Our results highlight a trade-off among speed, sensitivity, and accuracy of the ortholog search, with no single strategy performing optimally across all benchmark challenges. This underscores the need for flexibility in the strategy for orthology inference. fDA therefore provides the modular framework that allows users to tailor analyses to specific needs. However, for studies requiring accurate de novo gene structure prediction, we suggest using Augustus as the gene predictor.

This flexibility is particularly relevant considering existing approaches. Methods such as ALiBaSeq, Assexon, and eggNOG-mapper (28) are primarily designed for sequence retrieval and orthology assignment but not for comprehensive eukaryotic gene annotation. focus on sequence retrieval and orthology assignment rather than full eukaryotic gene annotation.

Although eggNOG-mapper can predict open reading frames in unannotated contigs, it relies on Prodigal (51), which was developed for prokaryotes and cannot model exon–intron structures. By contrast, BUSCO (23), Compleasm (29) and TOGA (27) perform explicit gene annotation. BUSCO and Compleasm use the same gene predictors as fDA but are limited to predefined BUSCO gene sets. TOGA projects annotations from a reference genome using whole-genome alignments, improving consistency across species but depending strongly on reference quality and potentially propagating annotation errors.

fDA fills a complementary niche by enabling targeted annotation and orthology inference for user-defined genes in unannotated genomes while also providing gene model quality assessment through feature architecture comparison.

### Performance of the orthology assignment

fDA builds on the targeted ortholog search tool fDOG (6). Our benchmarks show that both tools perform comparably to state-of-the-art software in identifying pairwise orthology relationships in comprehensive ortholog searches. (see Figs. 3 and 4, and (6)). The eggNOG mapper, in turn, which is mainly dedicated to the functional annotation of genomic and metagenomic data, provides only a quick and simple “first-pass” assignment of ortholog pairs (28). BUSCO and Compleasm use unidirectional profile-based searches in combination with predefined similarity thresholds for the ortholog identification. This strategy is admissible for single copy genes that have no extensive history of gene duplications or loss because no close paralogs can interfere with the ortholog assignment (see Fig. 3). However, for genes with more complex evolutionary histories including lineage-specific duplications and losses, the performances of both BUSCO and Compleasm are likely to significantly drop. Testing this is hard because BUSCO groups do not contain such genes. fDA can accurately infer orthologs even under such complex scenarios, although its sensitivity decreases compared to the analysis of single-copy genes.

The most comprehensive algorithm for ortholog searches in unannotated genomes is TOGA (27). However, whole genome alignments are computationally expensive, and thus it does not scale to the analysis of 10, 000+ genomes. Moreover, the use of TOGA is limited to taxa that are related closely enough such that also the non-coding regions parts in their genomes can be meaningfully aligned. Performing orthology assignments across phyla or kingdoms is therefore not possible. fDA, in contrast, performs similarity assessments on the amino acid level. Consequently, the evolutionary distances over which orthologs can be found depends only on the evolutionary rate of the proteins (45). fDA is therefore independent of evolutionary processes acting on non-coding parts of the genome and of its structure.

### Closing gaps in phylogenetic profiles

Genes that are overlooked during genome annotation result in gaps in phylogenetic profiles, thus generating a spurious signal of gene loss. Investigating the phylogenetic profiles of the BUSCO genes in the metazoan odb10 set revealed that ortholog searches using the annotated protein sets of 10 animal species resulted in 214 cases where an ortholog was missing in only 1 of the 10 analysed species (Supplementary Table S8). We hypothesized that at least a subset of these gaps is caused by annotation artefacts. Using targeted searches for orthologs in the unannotated genome sequences, we were able to fill more than 80 of these gaps in the phylogenetic profiles. In line with the artefact hypothesis, we found that mRNAs were mapped to the genomic locations where fDA identified an ortholog that was represented in the reference proteome of this species (see Supplementary Figure S12).

### Biosynthesis factors for beta-lactams in springtails

The presence of genes for β-lactam biosynthesis is common in springtails (see Fig. 5 and (47)) and beta-lactam production aligns well with the rich spectrum of other antimicrobial substances produced by these species (52). Our analysis has revealed that size and composition of enzyme repertoires vary among springtails.

At the lower end of the spectrum, we identify within the globular springtails (Symphypleona) a monophyletic clade that, with one exception, contains only taxa with an ortholog of pcbAB. Because the corresponding genome assemblies are highly fragmented (see Fig. 5E), orthologous loci for the other β-lactam biosynthetic genes could reside in assembly gaps.

However, most other assemblies analysed in this study are similarly fragmented, and the globular springtails containing only pcbAB do not show reduced BUSCO scores (see Fig. 5E). More importantly, orthologs of the remaining biosynthetic genes are absent from ten species in this clade that have been all sequenced and assembled independent from each other. Such a consistent pattern is difficult to explain if assembly gaps are randomly distributed across the genome. Instead, it is points towards a genuine absence of these β-lactam biosynthetic genes in this lineage. This observation also highlights the importance of broad taxon sampling, which is facilitated by fDA.

On the other end of the distribution, we find three species harbouring the entire gene cluster necessary for Cephalomycin C production, except for cmcH. This gene encodes a carbamoyltransferase, which adds a C-3’-hydroxyl group at the cephem nucleus (53). Its absence suggests that these species can probably synthesise a variant that differs slightly from Cephamycin C. Two of these species belong to the Onychiuridae (blind springtails) with a third member of this group, *Protaphorua quadriocellata*, also possessing orthologs to cefE and cefF, but not to the more common genes cmcI and cmcJ. Interestingly, we find further occurrences of cefE orthologs in other members in the clade harbouring the Onychiuridae but, with the exceptions of *C. granulata* and *D. trispinata*, not outside this clade. This suggests a clade-specific acquisition of these genes on the lineage leading to the blind springtails. Based on our findings, it will now be interesting to experimentally test the biosynthesis of these beta-lactams, to determine their precise molecular structure, to assess their physiological role and to what extent they contribute to the specific lifestyle of the blind springtails.

### Beta-lactam biosynthesis in other soil invertebrates

Thus far, little evidence existed for β-lactam synthesizing soil invertebrates outside the springtails (47). Here, we could identify orthologs to β-lactam biosynthesis genes in other taxa that had been sequenced in the context of the soil meta-invertebrate project (31). Among these, oribatid mites (Arachnida) are the second highest represented taxonomic group that is equipped at least with a basal gene set for β-lactam production (see Fig. 5). This parallels findings from the analysis of cellulose-degrading enzymes: these were also mainly found in springtails and in oribatid mites, but not in other soil invertebrates (12), probably indicating similarities in lifestyles. However, contrary to the findings for the cellulases, β-lactam synthesising genes are found only in a small fraction of the analysed oribatids. Also here, more complete genome assemblies will be required to assess whether this is an issue of data quality, or whether β-lactam biosynthesis is indeed only a sporadic trait among oribatids.

### The impact of assembly quality

Among all investigated enzymes involved into β-lactam biosynthesis, pcbAB is represented in the largest and taxonomically most diverse soil invertebrate species collection (see Fig. 5). In principle, this enzyme alone is sufficient for synthesizing the β-lactam ring (54). However, the feature architecture comparison between the reference protein from *Streptomyces clavuligerus* and of the individual orthologs, which is part of the fDA workflow, revealed a substantial heterogeneity. In some taxa, e.g., *H. dentifera* (Onychiuridae), fDA detected an ortholog whose feature architecture resembles that of the reference protein (see Fig. 5D). In others, e.g., *F. truncata*, the FAS score drops substantially, indicating that the ortholog shares no or only a single feature with the reference. While we cannot exclude a spurious orthology assignment of fDA, or a functional diversification of the ortholog, the likely explanation is different. pcbAB is a non-ribosomal peptide synthetase that extends over more than 3, 700 amino acids in *S. clavuligerus*. Its feature architecture is repetitive comprising, among others, four instances of a condensation domain (PF00668), three instances of an AMP binding domain (PF00501) and three instances of an AMP binding C domain (PF13193; Supplementary Fig. S14). fDA can faithfully annotate this gene in a genome assembly, which is indicated by the identification of orthologs with the same feature architecture as the reference protein. It is more likely that the assembly of the genomic region encoding such a repetitive protein itself is the problem (see (55).

### Relevance of beta-lactams

Our results, in agreement with Suring et al. (2017), indicate that β-lactam biosynthesis is a common trait in springtails, but may occur also in other soil invertebrates. Beside the relevance of these substances for the springtails, e.g., for keeping potentially harmful bacteria in rein, these findings have two further implications. First, soil invertebrates may serve as reservoir for complementing the portfolio of natural products available for developing novel antimicrobial substances (see (56)). Second, if springtails indeed produce antimicrobial substances in their gut, they might act as breeding factories for genes conveying antimicrobial resistance. Even though most environmental bacteria do not infect humans, they are capable of passing on their antimicrobial resistance genes to pathogenic relatives, potentially contributing to the emergence of multi- or even pan-drug resistant pathogens (57). Thus, our findings underline the contribution of the natural environment to the emergence of antibiotics resistances (58).

## Conclusions

In summary, fDOG-Assembly is the first targeted ortholog search tool that delivers the flexibility to search for orthologs to any gene of interest in un-annotated genome assemblies at a specificity that is comparable to that of protein-based ortholog search tools. A time and resource-consuming whole-genome annotation is by-passed by performing gene predictions in relevant genome positions on the fly. Additionally, the feature-architecture similarity between a seed gene and an identified ortholog is computed, which allows to trace likely changes in activity between orthologs over evolutionary time scales. We demonstrated the use of fDA by assessing the taxonomic distribution of beta-lactam biosynthesis capabilities across 176 soil invertebrates as an example application. This reveals that β-lactam biosynthesis appears far more prevalent than hitherto assumed in springtails, and that also individual oribatids are capable of β-lactam production. fDA therefore is a powerful yet flexible tool that significantly eases the access to high-resolution orthology-based evolutionary and functional studies independent of the annotation status of the underlying genomes.

## Supplementary Data

Supplementary Data are available at NAR Online

## Acknowledgements

The authors thank Felix Langschied for critically reading and discussing of the manuscript and the three anonymous reviewers for their constructive comments and suggestions during the review process.

