## Supplemental Figures and Tables for "Targeted ortholog search in unannotated genome assemblies with fDOG-Assembly"

### Supplementary material

**SUPPLEMENTARY MATERIAL** 1

**Supplementary Tables** 2
Table S1. Cross-reference table BUSCO ID to UniProt ID 2
Table S2. Precomputed Augustus models used for gene prediction with fDA-tb$_{Aug}$, fDA-MP$_{Aug}$ and BUSCO$_{Aug}$ 2
Table S3. Primer species used in the core ortholog compilation for 5,000 human proteins 2
Table S4. Primer taxa for compiling the core orthologous groups for the beta-lactam biosynthetic genes 3
Table S5. Proteins involved in beta-lactam biosynthesis used as seeds for the ortholog search with fDOG and fDOG-Assembly 4
Table S6. Ortholog predictions downloaded from the QfO benchmarking server 4
Table S7. Number of inferred orthologs per search species and corresponding benchmarking run 4
Table S8. Gaps in the phylogenetic profiles of the Metazoa odb10 dataset resolved per species 5
Table S9. Number of orthology assignments of fDA for set 2 6
Table S10. Performance of fDA in recovering human – Nematostella co-orthologous relationships predicted by InParanoid 6
Table S11. Soil invertebrate species used for beta-lactam gene search with fDA 6
Table S12. Assignment of soil invertebrate species to clusters of β-lactam biosynthesis repertoires 6
Table S13. Contamination screen of soil invertebrate orthologs to β-lactam biosynthesis genes 6
Table S14. Extended contamination screen for cefE and cefF considering genomic neighbourhood 6

**Supplementary Data** 7
Supplementary Data S1. Core orthologous groups used in the fDA runs 7
Supplementary Data S2. Phylogenetic profiles for beta-lactam biosynthesis factors 7

**Supplementary Figures** 8
Supplementary Figure S1. Mapping of UniProt-IDs to fDA gene predictions. 8
Supplementary Figure S2. Agreement between the BUSCO$_{Aug}$ gene annotation and the pre-annotated genes in the individual QfO reference species. ‘ 9
Supplementary Figure S3. Agreement between the BUSCO$_{ME}$ gene annotation and the pre-annotated genes in the individual QfO reference species. 9
Supplementary Figure S4. Agreement between the fDA-tb$_{Aug}$ gene annotation and the pre-annotated genes in the individual QfO reference species. 10
Supplementary Figure S5. Agreement between the fDA-tb$_{ME}$ gene annotation and the pre-annotated genes in the individual QfO reference species. 10
Supplementary Figure S6. Agreement between the fDA-MP gene annotation and the pre-annotated genes in the individual QfO reference species. 11
Supplementary Figure S7. Agreement between the fDA-MP$_{Aug}$ gene annotation and the pre-annotated genes in the individual QfO reference species. 11
Supplementary Figure S8. Agreement between the fDA-MP$_{ME}$ gene annotation and the pre-annotated genes in the individual QfO reference species. 12
Supplementary Figure S9. Agreement between the Compleasm gene annotation and the pre-annotated genes in the individual QfO reference species. 12
Supplementary Figure S10. Overlap of the ortholog assignments made by fDA, Compleasm and BUSCO. 13
Supplementary Figure S11. Consistency of orthology assignments from assembly-based tools with those from proteome-based tools. 13
Supplementary Figure S12. The fDA-tb$_{AUG}$ ortholog to the BUSCO gene 613851at33208 overlaps with a curated NCBI RefSeq gene that is not included in the QfO reference proteome. 13
Supplementary Figure S13. Phylogenetic Profiles of genes involved in beta-lactam biosynthesis across soil-living invertebrates. 14
Supplementary Figure S14. Domain Architecture of pcbAB (d-(L-a-Aminoadipyl)-L-Cysteinyl-D-Valine Synthetase) from Streptomyces clavuligerus. 14

#### Supplementary Tables

##### Table S1. Cross-reference table BUSCO ID to UniProt ID

See *separate file Supplementary_Table_1-Muelbaier-fDA.xlsx*

##### Table S2. Precomputed Augustus models used for gene prediction with fDA-tb$_{Aug}$, fDA-MP$_{Aug}$ and BUSCO$_{Aug}$

| Genome assembly | Augustus model |
|---|---|
| *Caenorhabditis elegans* | caenorhabditis |
| *Danio rerio* | zebrafish |
| *Drosophila melanogaster* | fly |
| *Gallus gallus* | chicken |
| *Helobdella robusta* | caenorhabditis |
| *Ixodes scapularis* | fly |
| *Nematostella vectensis* | fly |
| *Rattus norvegicus* | human |
| *Tribolium castaneum* | tribolium2012 |
| *Xenopus tropicalis* | human |

##### Table S3. Primer species used in the core ortholog compilation for 5,000 human proteins

| UniprotID | Species |
|---|---|
| UP000007062 | *Anopheles gambiae* (African malaria mosquito) |
| UP000000589 | *Mus musculus* (Mouse) |
| UP000005640 | *Homo sapiens* (Human) |
| UP000009136 | *Bos taurus* (Bovine) |
| UP000001038 | *Oryzias latipes* (Japanese rice fish) |
| UP000002280 | *Monodelphis domestica* (Gray short-tailed opossum) |
| UP000001519 | *Gorilla gorilla gorilla* (Western lowland gorilla) |
| UP000002254 | *Canis lupus familiaris* (Dog) |
| UP000002277 | *Pan troglodytes* (Chimpanzee) |
| UP000001554 | *Branchiostoma floridae* (Florida lancelet) |
| UP000018468 | *Lepisosteus oculatus* (Spotted gar) |

**Table S4. Primer taxa for compiling the core orthologous groups for the beta-lactam biosynthetic genes**

| Species | NCBI ID | RefSeq ID | Identifier |
|---|---|---|---|
| *Amycolatopsis albispora* | 1804986 | GCF_003312875.1 | AMYAL@1804986@003312875_1 |
| *Amycolatopsis magusensis* | 882444 | GCF_017875555.1 | AMYMA@882444@017875555_1 |
| *Aspergillus nidulans* | 227321 | GCF_000149205.2 | ASPNI@227321@000149205_2 |
| *Aspergillus puulaauensis* | 1220207 | GCF_016861865.1 | ASPPU@1220207@016861865_1 |
| *Hyaloscypha bicolor* | 1095630 | GCF_002865645.1 | HYABI@1095630@002865645_1 |
| *Modestobacter excelsi* | 2213161 | GCF_005930495.1 | MODEX@2213161@005930495_1 |
| *Streptantibioticus cattleyicolor* | 1003195 | GCF_000237305.1 | STRCA@1003195@000237305_1 |
| *Streptomyces clavuligerus* | 1901 | GCF_000148465.1 | STRCL@1901@000148465_1 |
| *Streptomyces jumonjinensis* | 1945 | GCF_009600885.1 | STRJU@1945@009600885_1 |
| *Streptomyces katsurahamanus* | 2577098 | GCF_009600895.1 | STRKA@2577098@009600895_1 |
| *Streptomyces megasporus* | 44060 | GCF_000718985.1 | STRME@44060@000718985_1 |
| *Streptomyces odonnellii* | 1417980 | GCF_000981895.1 | STROD@1417980@000981895_1 |
| *Streptomyces sulfonofaciens* | 68272 | GCF_014656295.1 | STRSU@68272@014656295_1 |
| *Streptomyces tirandamycinicus* | 2174846 | GCF_003097515.1 | STRTI@2174846@003097515_1 |
| *Trichophyton benhamiae* | 663331 | GCF_000151125.1 | TRIBE@663331@000151125_1 |
| *Trichophyton rubrum* | 559305 | GCF_000151425.1 | TRIRU@559305@000151425_1 |
| *Trichophyton verrucosum* | 663202 | GCF_000151505.1 | TRIVE@663202@000151505_1 |

**Table S5. Proteins involved in beta-lactam biosynthesis used as seeds for the ortholog search with fDOG and fDOG-Assembly**

| Gene Name | Reference species | NCBI taxonomy ID | Uniprot ID | NCBI Accession |
|---|---|---|---|---|
| **cefE** | *Streptomyces clavuligerus* | 1901 | P18548 | WP_003952493.1 |
| **cefF** | *Streptomyces clavuligerus* | 1901 | P42220 | WP_003952498.1 |
| **cmcH** | *Streptomyces clavuligerus* | 1901 | O85728 | WP_003952499.1 |
| **cmcI** | *Streptomyces clavuligerus* | 1901 | B5GLB3 | WP_003952496.1 |
| **cmcJ** | *Streptomyces clavuligerus* | 1901 | B5GLB4 | WP_003952497.1 |
| **cefD** | *Streptomyces clavuligerus* | 1901 | P18549 | WP_003952494.1 |
| **pcbC** | *Streptomyces clavuligerus* | 1901 | P10621 | WP_003952506.1 |
| **pcbAB** | *Streptomyces clavuligerus* | 1901 | Q01757 | WP_003952505.1 |
| **penDE** | *Aspergillus nidulans* | 162425 | P21133 | XP_660227.1 |

**Table S6. Ortholog predictions downloaded from the QfO benchmarking server**

| Dataset |
|---|
| BBH |
| Domainoid+ |
| EnsemblCompara |
| Hieranoid2 |
| InParanoid5 |
| MethaPhOrs v2.5 |
| OMA pairs |
| OrthoFFGC |
| OrthoFinder 2.5.5 |
| OrthoInspector 3.5 |
| PANTHER 18.0 all |
| RSD |
| SonicParanoid sens |

**Table S7. Number of inferred orthologs per search species and corresponding benchmarking run**

| | fDA-tb | | fDA-MP | | | BUSCO | | Compleasm |
|---|---|---|---|---|---|---|---|---|
| | Aug | ME | Aug | ME | MP | Aug | ME | MP |

| | | | | | | | | |
|---|---|---|---|---|---|---|---|---|
| *C. elegans* | 728 | 735 | 690 | 636 | 646 | 741 | 739 | 729 |
| *D. rerio* | 1164 | 1169 | 1158 | 1163 | 1165 | 1245 | 1156 | 1155 |
| *D. melanogaster* | 926 | 940 | 958 | 958 | 966 | 960 | 961 | 957 |
| *G. gallus* | 897 | 918 | 909 | 906 | 908 | 914 | 917 | 918 |
| *H. robusta* | 803 | 870 | 915 | 947 | 939 | 895 | 972 | 921 |
| *I. scapularis* | 863 | 971 | 961 | 1049 | 982 | 894 | 919 | 928 |
| *N. vectensis* | 872 | 1049 | 944 | 1019 | 1007 | 935 | 957 | 948 |
| *R. norvegicus* | 1009 | 1073 | 1015 | 1029 | 1045 | 1020 | 1276 | 1060 |
| *T. castaneum* | 945 | 952 | 955 | 967 | 961 | 959 | 960 | 955 |
| *X. tropicalis* | 949 | 975 | 952 | 955 | 952 | 958 | 958 | 958 |
| **Total** | **9156** | **9652** | **9457** | **9629** | **9571** | **9521** | **9815** | **9529** |

**Table S8. Gaps in the phylogenetic profiles of the Metazoa odb10 dataset resolved per species**

| | **Gaps$** |
|---|---|
| *Gallus gallus* | 32 |
| *Rattus norvegicus* | 5 |
| *Tribolium castaneum* | 4 |
| *Caenorhabditis elegans* | 55 |
| *Nematostella vectensis* | 26 |
| *Ixodes scapularis* | 43 |
| *Danio rerio* | 1 |
| *Helobdella robusta* | 21 |
| *Xenopus tropicalis* | 14 |
| *Drosophila melanogaster* | 13 |
| **Total** | **214** |

$ We only consider gaps where 9 of the 10 species are represented by an ortholog

**Table S9. Number of orthology assignments of fDA for set 2**

| | Rat | *N. vectensis* | Total |
|---|---|---|---|
| | #orthologs (#mapped to UniProt) | #orthologs (#mapped to UniProt) | #orthologs (#mapped to UniProt) |
| **fDA-tb$_{Aug}$** | 5,749 (4,920) | 3,246 (2,702) | 8,995 (7,622) |
| **fDA-tb$_{ME}$** | 5,642 (4,781) | 4,365 (3,441) | 10,007 (8,222) |
| **fDA-MP$_{Aug}$** | 5,104 (4,653) | 2,318 (1,941) | 9,468 (6,594) |
| **fDA-MP$_{ME}$** | 5,442 (4,634) | 2,821 (2,190) | 8,263 (6,824) |
| **fDA-MP** | 5,021 (4,703) | 2,434 (2,119) | 7,455 (6,822) |

**Table S10. Performance of fDA in recovering human – Nematostella co-orthologous relationships predicted by InParanoid**

| **Group layout** | | **1:1** | **1:n** | **n:1** | **n:n** |
|---|---|---|---|---|---|
| | | **Orthology assignments: #groups (#pairs)** | | | |
| *N. vectensis* | InParanoid | 1425 (1425) | 167 (527) | 145 (334) | 59 (1280) |
| | fDA-tb$_{Aug}$ | 762 (762) | 76 (115) | 84 (156) | 24 (114) |
| | fDA-tb$_{ME}$ | 1117 (1117) | 105 (200) | 107 (202) | 29 (143) |
| | fDA-MP$_{Aug}$ | 797 (797) | 73 (89) | 95 (181) | 21 (56) |
| | fDA-MP$_{ME}$ | 1032 (1032) | 90 (127) | 113 (221) | 22 (62) |
| | fDA-MP | 1023 (1023) | 91 (224) | 113 (123) | 23 (75) |

**Table S11. Soil invertebrate species used for beta-lactam gene search with fDA**

*See Supplementary_Table_11_Muelbaier_fDA.xlsx*

**Table S12. Assignment of soil invertebrate species to clusters of β-lactam biosynthesis repertoires**

*See Supplementary_Table_12_Muelbaier_fDA.xlsx*

**Table S13. Contamination screen of soil invertebrate orthologs to β-lactam biosynthesis genes**

*See Supplementary_Table_13_Muelbaier_fDA.xlsx*

**Table S14. Extended contamination screen for cefE and cefF considering genomic neighbourhood**

*See Supplementary_Table_14_Muelbaier_fDA.xlsx*

#### Supplementary Data

**Supplementary Data S1. Core orthologous groups used in the fDA runs**

*See Supplementary_Data_1_core-orthologous-groups.tar.gz*

**Supplementary Data S2. Phylogenetic profiles for beta-lactam biosynthesis factors**

*See Supplementary_Data_2_fDA-output.tar.gz*

#### Supplementary Figures

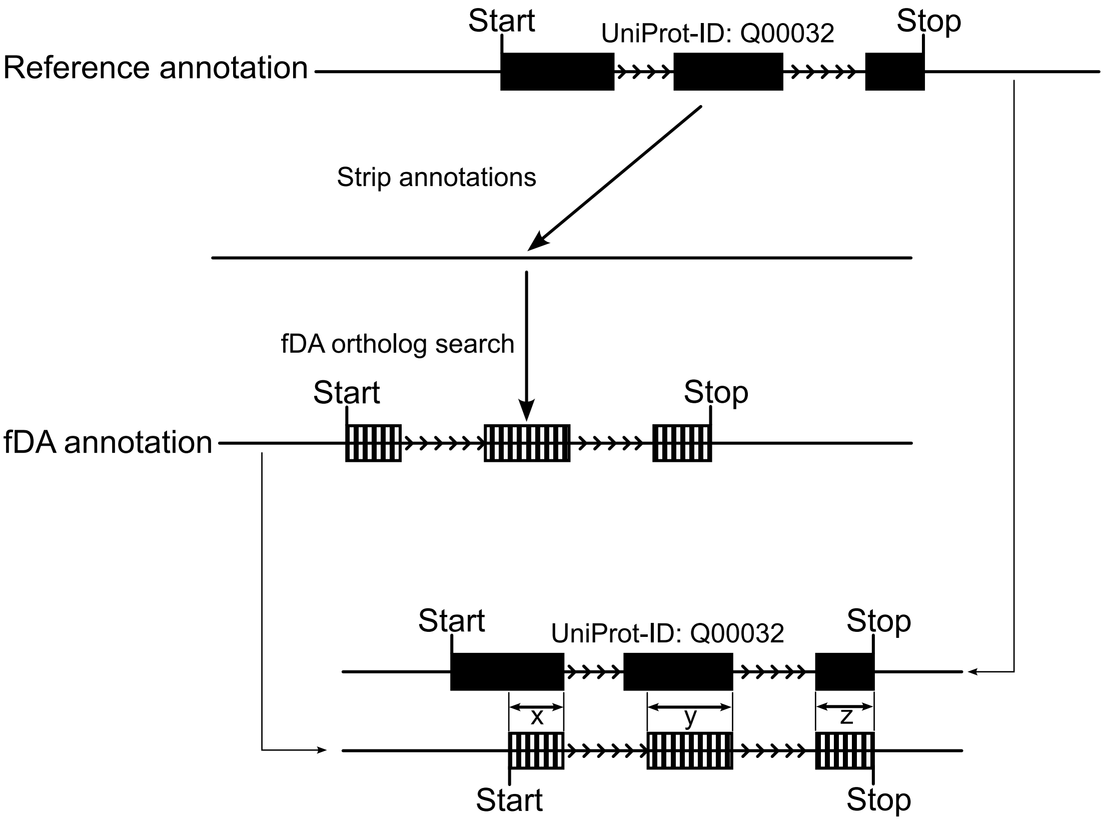


**Supplementary Figure S1. Mapping of UniProt-IDs to fDA gene predictions.** Reference gene annotations are stripped from the assembly. The assembly is then used for an fDA ortholog search. To assign the ortholog identified by fDA a UniProt-ID, we compare the genomic positions of the corresponding exons to those in the reference annotation and determine the overlap (x, y, and z in the figure). If the two gene annotations reside on the same strand, and if the sum of the overlaps exceeds 0.5 x length of the reference gene, we transfer the UniProt-ID of the corresponding protein to that encoded by the fDA annotated gene. See Supplementary Figs. S2 – S8 for a distribution of overlaps for the different ortholog search tools and search modes.

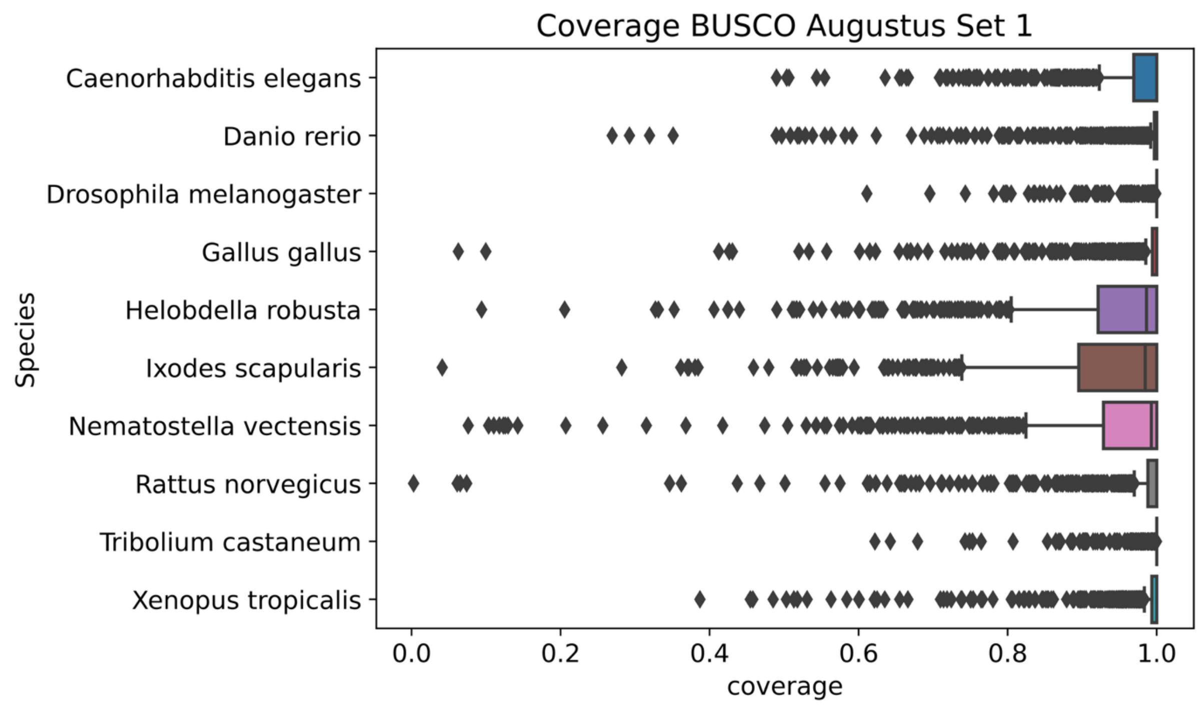


**Supplementary Figure S2. Agreement between the BUSCO$_{Aug}$ gene annotation and the pre-annotated genes in the individual QfO reference species.** ‘Coverage’ denotes the fraction of nucleotides in the protein-coding exons of the pre-annotated genes that are contained in protein-coding exons of the BUSCO-annotated genes (see Supplementary Figure S1 for details). The coverage distribution is provided for the genes in benchmark set 1.

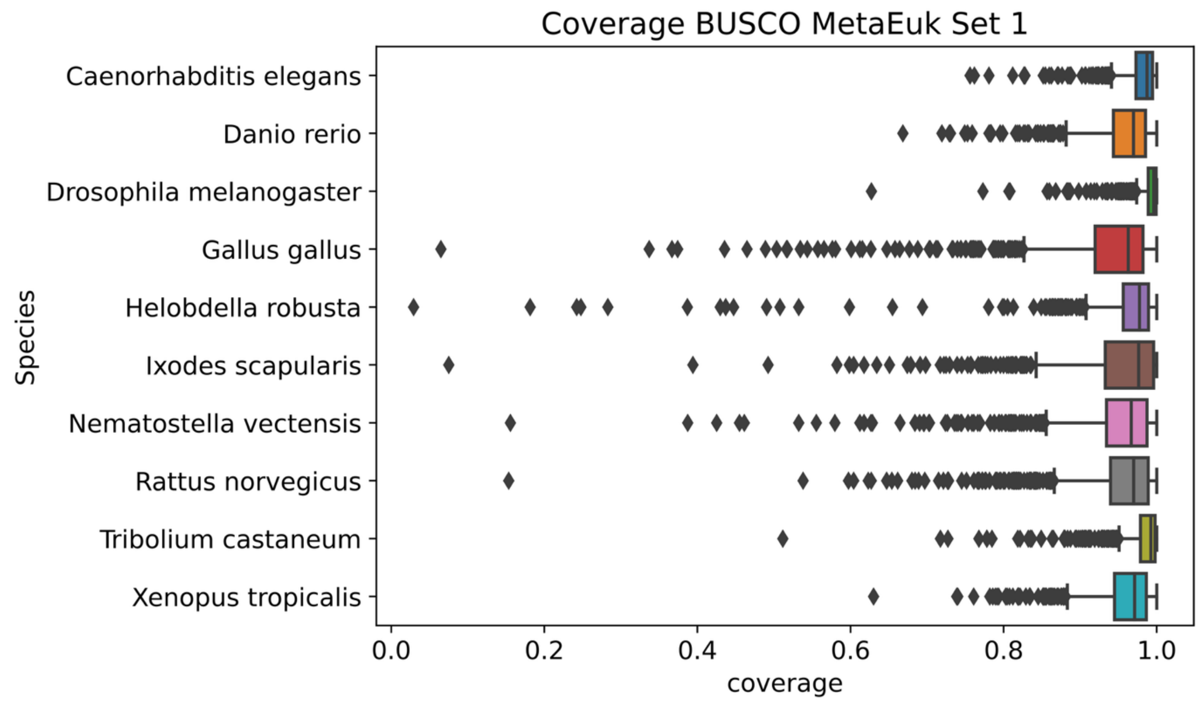


**Supplementary Figure S3. Agreement between the BUSCO$_{ME}$ gene annotation and the pre-annotated genes in the individual QfO reference species.** ‘Coverage’ denotes the fraction of nucleotides in the protein-coding exons of the pre-annotated genes that are contained in protein-coding exons of the BUSCO$_{ME}$-annotated genes (see Supplementary Figure S1 for details). The coverage distribution is provided for the genes in benchmark set 1.

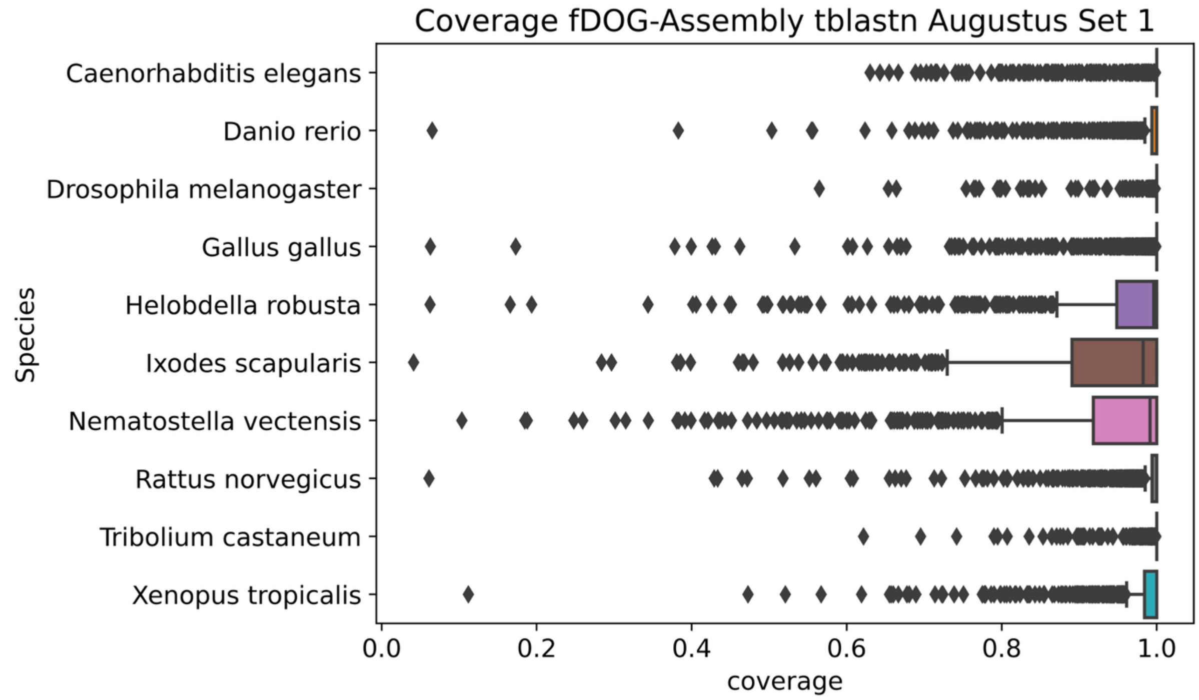


**Supplementary Figure S4. Agreement between the fDA-tb$_{Aug}$ gene annotation and the pre-annotated genes in the individual QfO reference species.** 'Coverage' denotes the fraction of nucleotides in the protein-coding exons of the pre-annotated genes that are contained in protein-coding exons of the fDA-tb$_{Aug}$-annotated genes (see Supplementary Figure S1 for details). The coverage distribution is provided for the genes in benchmark set 1.

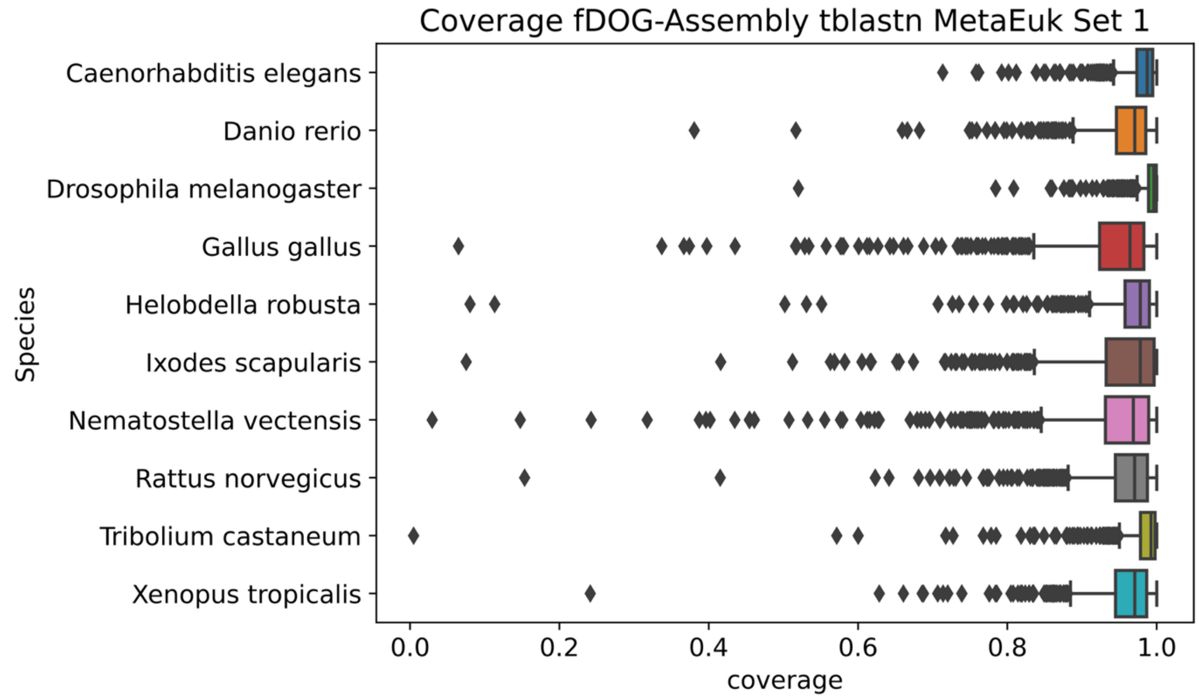


**Supplementary Figure S5. Agreement between the fDA-tb$_{ME}$ gene annotation and the pre-annotated genes in the individual QfO reference species.** 'Coverage' denotes the fraction of nucleotides in the protein-coding exons of the pre-annotated genes that are contained in protein-coding exons of the fDA-tb$_{ME}$-annotated genes (see Supplementary Figure S1 for details). The coverage distribution is provided for the genes in benchmark set 1.

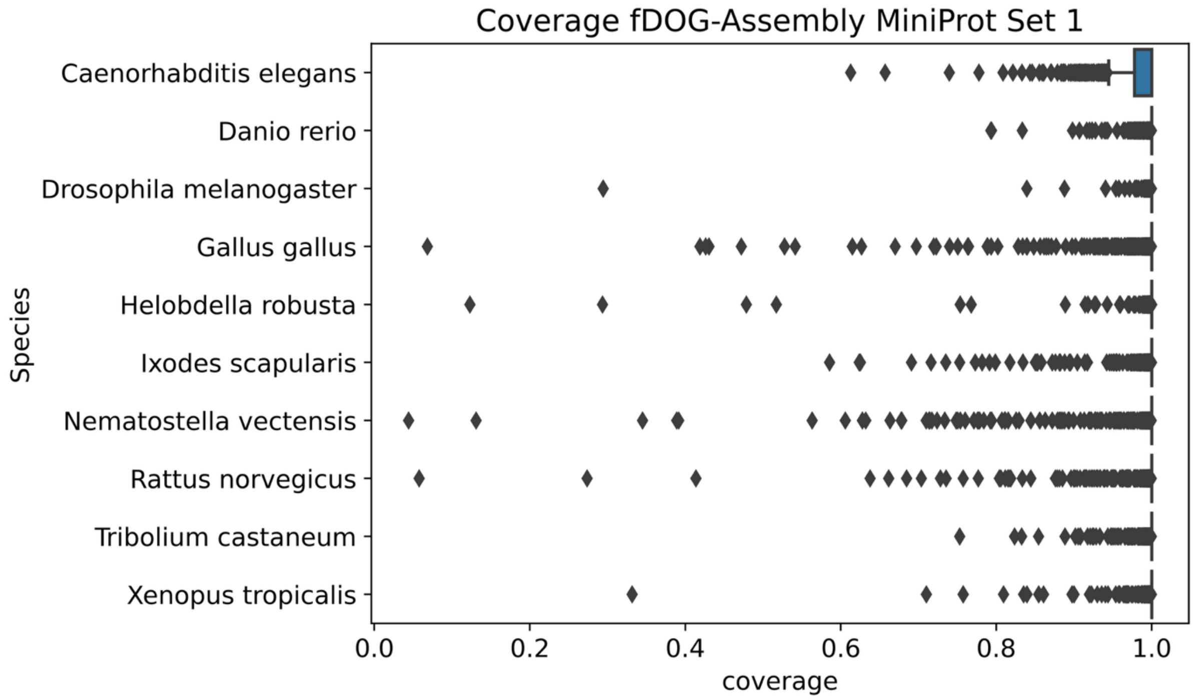


**Supplementary Figure S6. Agreement between the fDA-MP gene annotation and the pre-annotated genes in the individual QfO reference species.** 'Coverage' denotes the fraction of nucleotides in the protein-coding exons of the pre-annotated genes that are contained in protein-coding exons of the fDA-MP-annotated genes (see Supplementary Figure S1 for details). The coverage distribution is provided for the genes in benchmark set 1.

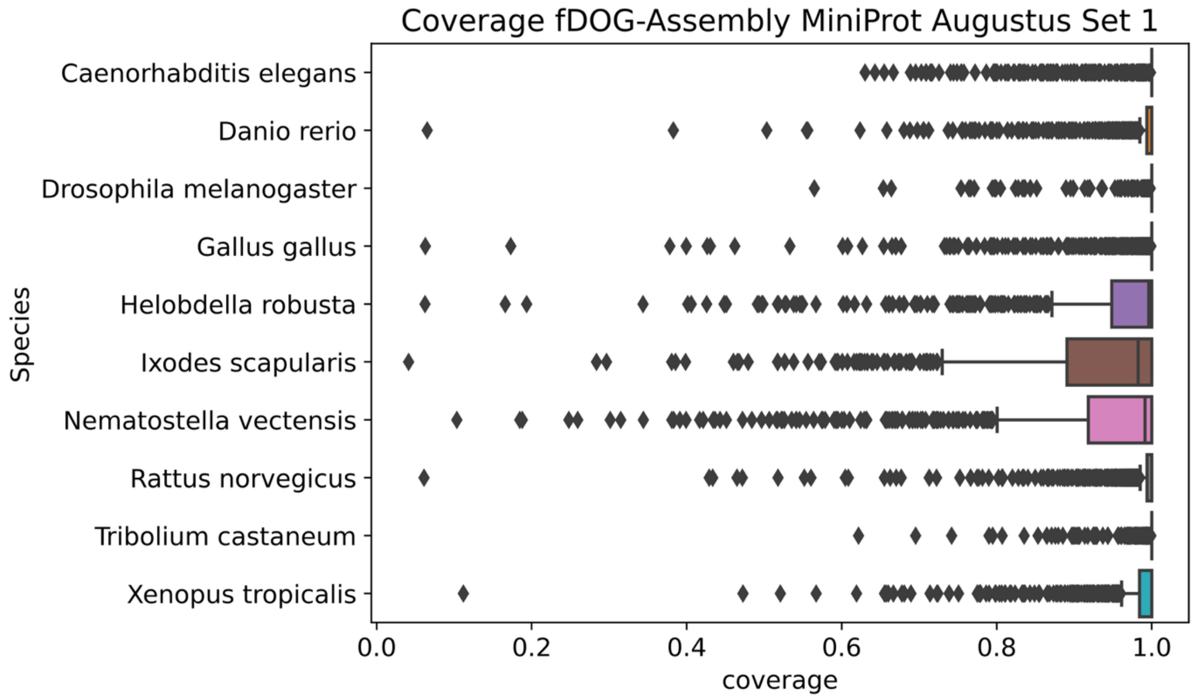


**Supplementary Figure S7. Agreement between the fDA-MP$_{Aug}$ gene annotation and the pre-annotated genes in the individual QfO reference species.** 'Coverage' denotes the fraction of nucleotides in the protein-coding exons of the pre-annotated genes that are contained in protein-coding exons of the fDA-MP$_{Aug}$-annotated genes (see Supplementary Figure S1 for details). The coverage distribution is provided for the genes in benchmark set 1.

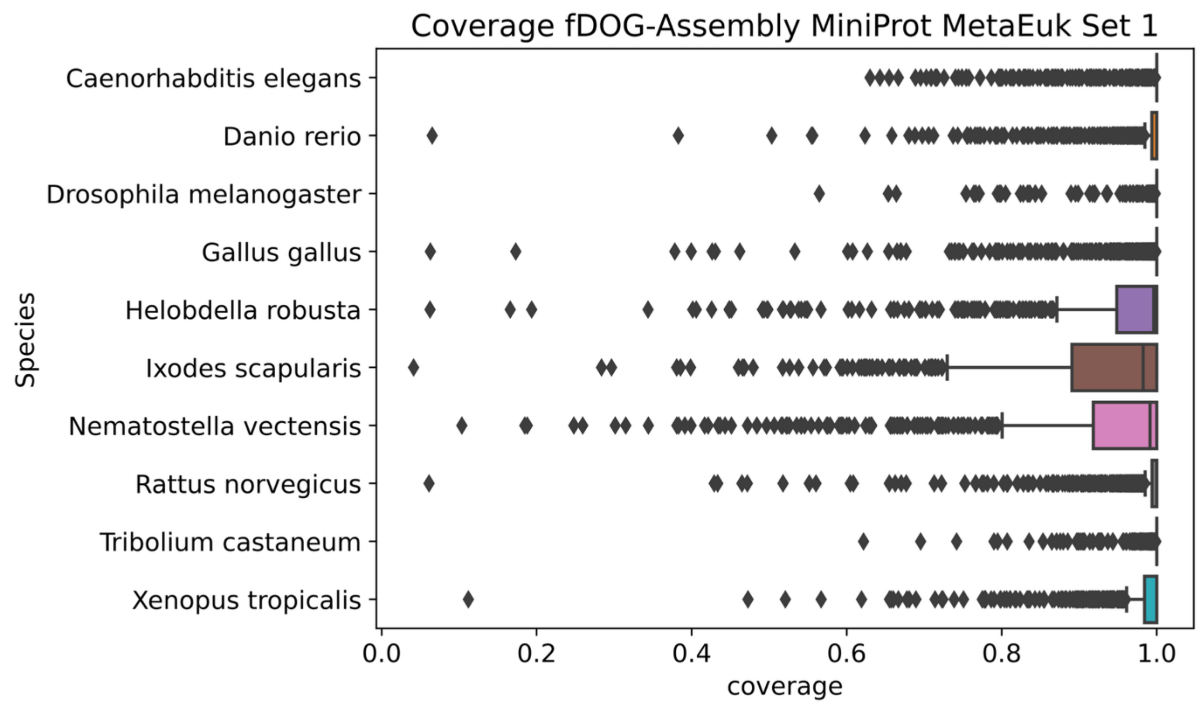


**Supplementary Figure S8. Agreement between the fDA-MP$_{ME}$ gene annotation and the pre-annotated genes in the individual QfO reference species.** 'Coverage' denotes the fraction of nucleotides in the protein-coding exons of the pre-annotated genes that are contained in protein-coding exons of the fDA-MP$_{ME}$-annotated genes (see Supplementary Figure S1 for details). The coverage distribution is provided for the genes in benchmark set 1.

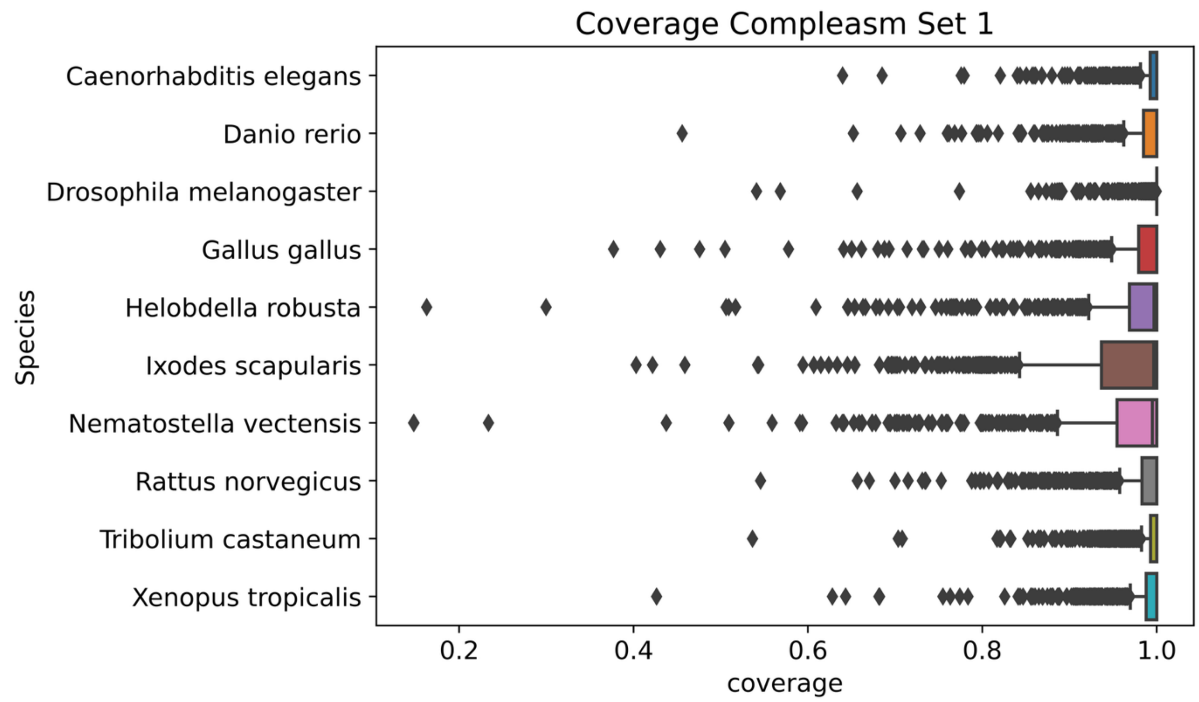


**Supplementary Figure S9. Agreement between the Compleasm gene annotation and the pre-annotated genes in the individual QfO reference species.** 'Coverage' denotes the fraction of nucleotides in the protein-coding exons of the pre-annotated genes that are contained in protein-coding exons of the Compleasm-annotated genes (see Supplementary Figure S1 for details). The coverage distribution is provided for the genes in benchmark set 1.

**Supplementary Figure S10. Overlap of the ortholog assignments made by fDA, Compleasm and BUSCO.** The number of orthologs identified in the individual searches is provided in the first column and indicated by the bar. Patterns of stacked filled circles indicate orthologs consistently identified by the respective tools, and the total number per pattern is given in the bar chart. See *Supplementary_Figure_S10_Muelbaier_fDA.pdf*

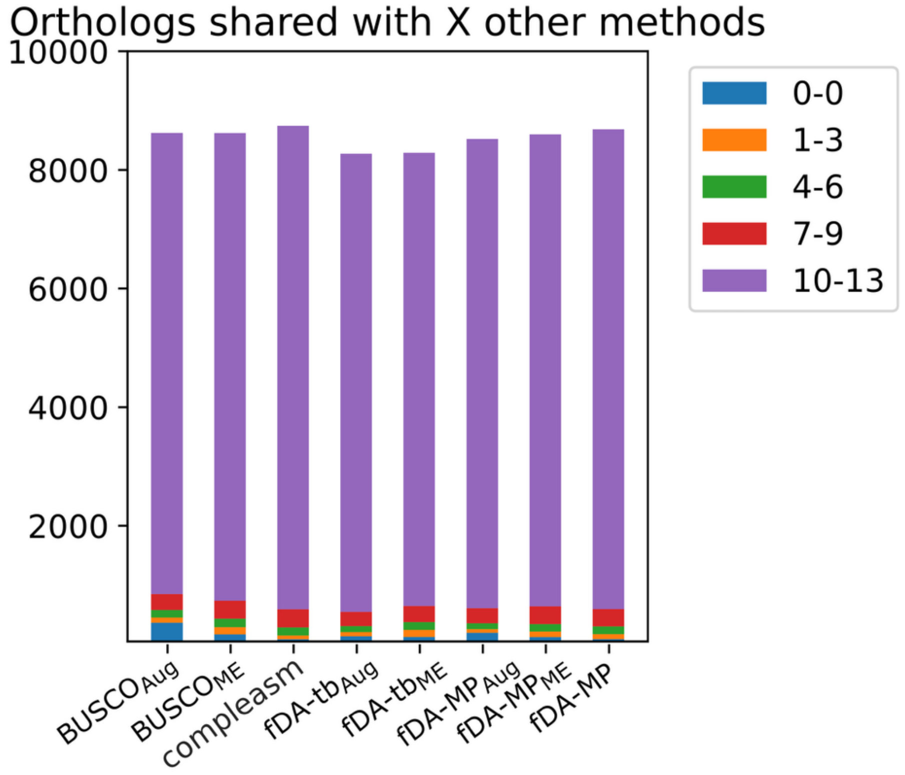


**Supplementary Figure S11. Consistency of orthology assignments from assembly-based tools with those from proteome-based tools.**
The color code indicates how many of the proteome-based tools support each assignment made by the respective assembly-based tool. Bar heights specify the number of ortholog assignments within each category (linear scale).

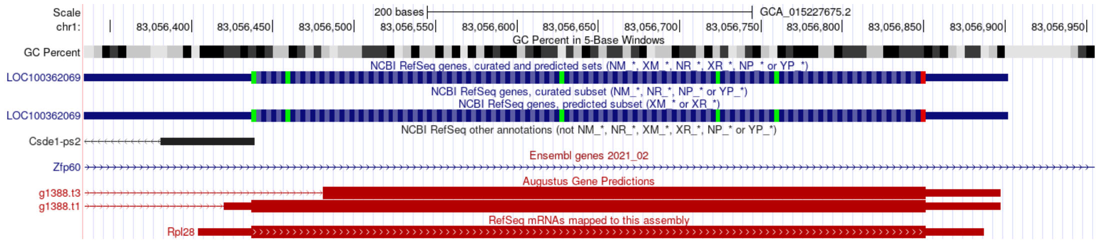


**Supplementary Figure S12. The fDA-tb$_{AUG}$ ortholog to the BUSCO gene 613851at33208 overlaps with a curated NCBI RefSeq gene that is not included in the QfO reference proteome.** fDA$_{Aug}$ identified an ortholog in *Rattus norvegicus* (GCA_015227675) that closes a gap in the phylogenetic profile that was based on the analysis of the QfO reference proteomes using conventional ortholog search tools. Mapping the identified ortholog to the genome assembly (Augustus gene prediction) reveals that it fully overlaps with an annotated RefSeq protein and a RefSeq mRNA.

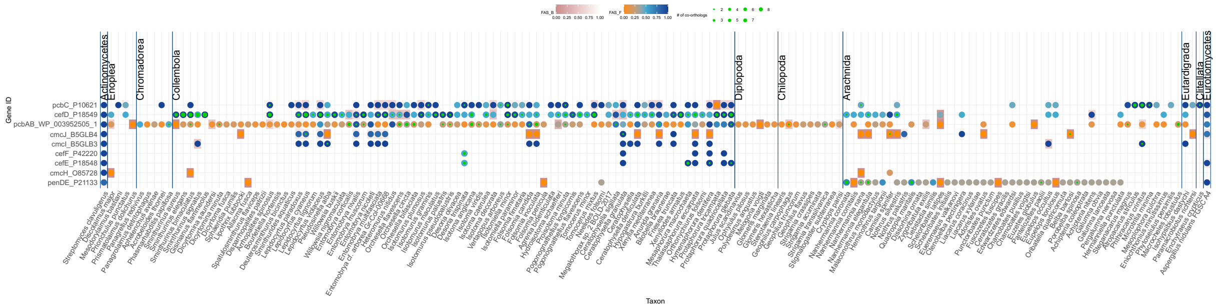

**Supplementary Figure S13. Phylogenetic Profiles of genes involved in beta-lactam biosynthesis across soil-living invertebrates.** Each row represents the phylogenetic profile for the gene indicated in the first column. The first and the last column represent *Streptomyces clavuligerus*, and *Aspergillus nidulans*, respectively. *S. clavuligerus* served as the reference species for all fDA searches except for penDE, where *A. nidulans* was used. Dots indicate the detection of an ortholog to the respective seed gene in the individual species. The dot color encodes the feature architecture similarity between the seed protein and its ortholog using the seed architecture as reference. The color gradient ranges from blue (high similarity) to orange (low similarity). The cell color encodes the feature architecture similarity between the seed protein and its ortholog using the ortholog architecture as reference. The color gradient ranges from white (high similarity) to pink (low similarity). Green inlays indicate that fDA detected more than one co-ortholog.

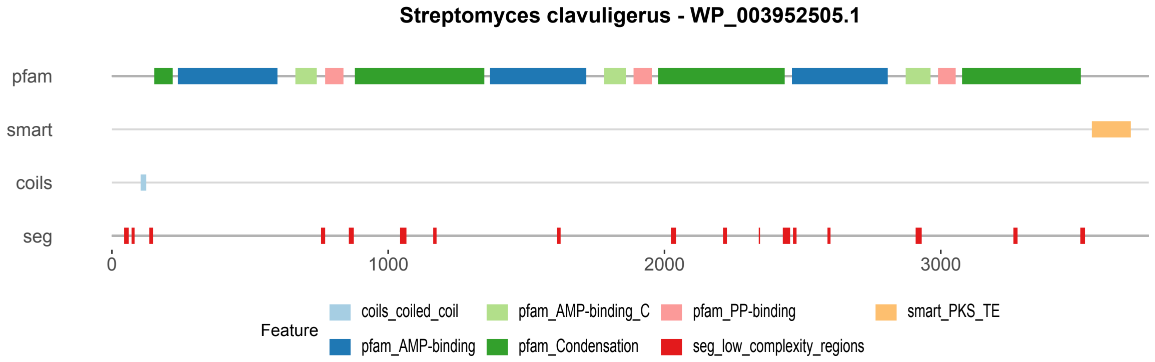


**Supplementary Figure S14. Domain Architecture of pcbAB (d-(L-a-Aminoadipyl)-L-Cysteinyl-D-Valine Synthetase) from Streptomyces clavuligerus.** Pfam_AMP-binding - PF00501; pfam_AMP-binding_C - PF13193; pfam_Condensation - PF00668; pfam_PP-binding - PF00550; smart_PKS_TE - SM000824
