## Supplementary figures and images for "Targeted ortholog search in unannotated genome assemblies with fDOG-Assembly"

### Supplemental Fig. S10

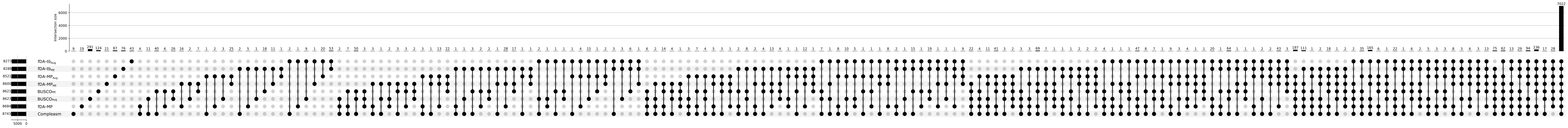
